# Identification of a new family of peptidoglycan transpeptidases reveals atypical crosslinking is essential for viability in *Clostridioides difficile*

**DOI:** 10.1101/2024.03.14.584917

**Authors:** Kevin W. Bollinger, Ute Müh, Karl L. Ocius, Alexis J. Apostolos, Marcos M. Pires, Richard F. Helm, David L. Popham, David S. Weiss, Craig D. Ellermeier

**Affiliations:** Department of Microbiology and Immunology Carver College of Medicine, University of Iowa Iowa City, IA, USA; Department of Chemistry University of Virginia Charlottesville, VA, USA; Department of Biochemistry Virginia Tech, Blacksburg, VA, USA; Department of Biological Sciences Virginia Tech, Blacksburg, VA, USA; Graduate Program in Genetics University of Iowa, Iowa City, IA USA; Haleon, 1211 Sherwood Ave, Richmond, VA 23220

## Abstract

*Clostridioides difficile*, the leading cause of antibiotic-associated diarrhea, relies primarily on 3-3 crosslinks created by L,D-transpeptidases (LDTs) to fortify its peptidoglycan (PG) cell wall. This is unusual, as in most bacteria the vast majority of PG crosslinks are 4-3 crosslinks, which are created by penicillin-binding proteins (PBPs). Here we report the unprecedented observation that 3-3 crosslinking is essential for viability in *C. difficile*. We also report the discovery of a new family of LDTs that use a VanW domain to catalyze 3-3 crosslinking rather than a YkuD domain as in all previously known LDTs. Bioinformatic analyses indicate VanW domain LDTs are less common than YkuD domain LDTs and are largely restricted to Gram-positive bacteria. Our findings suggest that LDTs might be exploited as targets for antibiotics that kill *C. difficile* without disrupting the intestinal microbiota that is important for keeping *C. difficile* in check.

## INTRODUCTION

*Clostridioides difficile* is a Gram-positive, spore-forming opportunistic pathogen that has become the leading cause of antibiotic-associated diarrhea in high-income countries. The CDC estimates that *C. difficile* infections kill over 12,000 people per year in the United States^1^. *C. difficile* infections are often triggered by broad-spectrum antibiotics administered either prophylactically or to treat some other infection. These antibiotics have the unintended consequence of disrupting the intestinal microbiota that ordinarily keeps *C. difficile* in check^2,3^. The frontline treatment for *C. difficile* infections is vancomycin, which is usually effective but also kills desirable bacteria, so relapse rates exceed 20%, and for this cohort the prognosis is poor^4–6^. An antibiotic that kills *C. difficile* more selectively would presumably improve outcomes, but developing such a drug requires identifying targets uniquely important to *C. difficile*.

Many of our most useful antibiotics target biogenesis of the bacterial cell wall, which provides essential protection against lysis due to turgor pressure. The cell wall is composed of peptidoglycan (PG), a complex meshwork of glycan strands of alternating *N*-acetylglucosamine (NAG) and *N*-acetylmuramic acid (NAM) that are stitched together by short peptide crosslinks^7,8^. The predominant modes of crosslinking, termed 4-3 and 3-3, are named based on the position of the amino acids involved; 4-3 crosslinks join a D-Alanine (D-Ala) in position four of one peptide to a *meso*-diaminopimelic acid (*m*DAP) in position three of another, while 3-3 crosslinks join two *m*DAP residues (Fig 1a). In most well-studied bacteria, as exemplified by *Escherichia coli*, about 90% of the crosslinks are 4-3, and these are essential for viability, whereas only ∼10% of the crosslinks are 3-3 and these are not essential ^9,10^.

**Fig 1.**
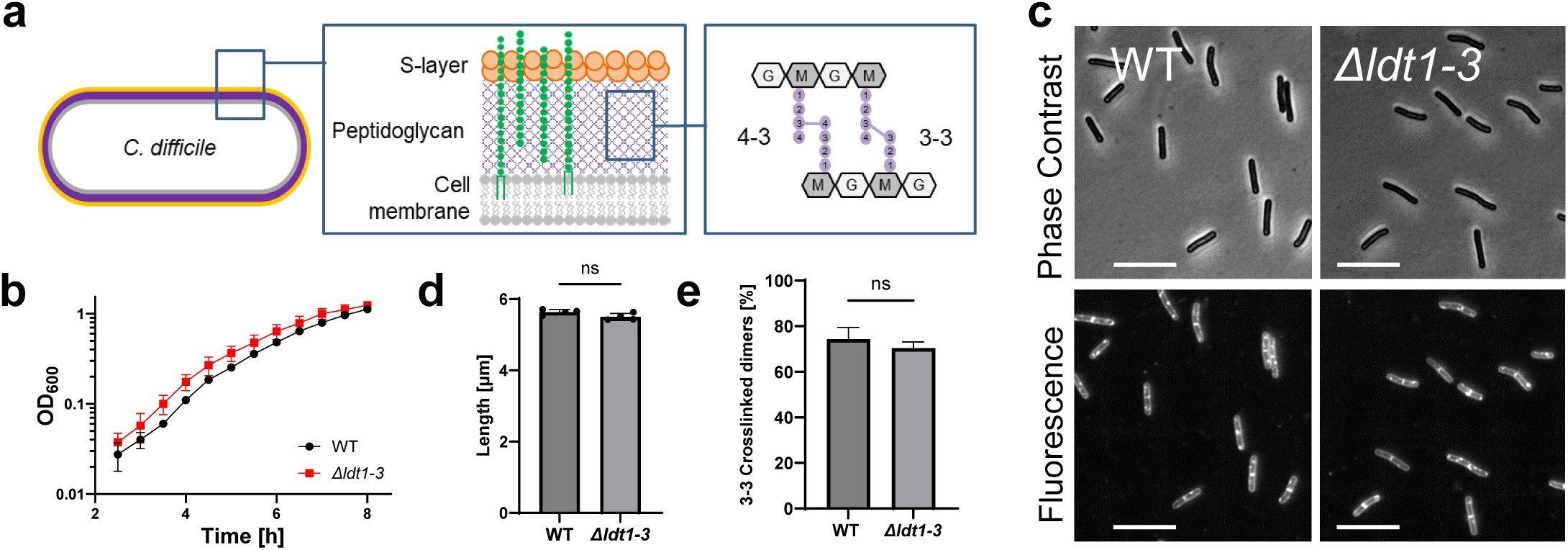
A *C. difficile* mutant lacking all three YkuD-type Ldts (*Δldt1-3*) exhibits wild-type growth, morphology and 3-3 crosslinking. (**a**) Diagram of the cell envelope of *C. difficile*. The PG matrix contains a repeating disaccharide of *N*-acetylglucosamine (NAG, G) and *N*-acetylmuramic acid (NAM, M). The glycans are crosslinked by short peptides (filled circles) attached to the NAM residues. About 25% of the crosslinks are 4-3 crosslinks created by PBPs and 75% are 3-3 crosslinks created by LDTs. Polysaccharides (green) analogous to teichoic acids are attached to PG or to a lipid in the cell membrane. (**b**) Growth curve in TY. Filled symbols and error bars indicate the mean ± s.d. from four biological replicates. (**c**) Phase contrast and fluorescence micrographs of cells sampled at OD_600_ = 0.5 and stained with the membrane dye FM4-64. Size bars, 10 μm. Images representative of 3 experiments. (**d**) Average cell length based on four biological replicates in which >160 cells were measured per sample. Dots depict the mean value from each sample, bars and error bars the mean ± s.d. across all four trials. ns, not significant in an unpaired two-tailed *t*-test. (**e**) Percentage of 3-3 PG crosslinks as a fraction of the total crosslinked dimers graphed as mean ± s.d. from three biological replicates. ns, not significant in an unpaired one-tailed *t*-test. Strains used: WT = R20291, Δ*ldt1-3* = KB124

The two modes of crosslinking rely on completely different enzymes. All 4-3 crosslinks are synthesized by D,D-transpeptidases, more commonly referred to as penicillin-binding proteins (PBPs)^11^. As the name suggests, PBPs are the lethal targets of penicillin and other β-lactam antibiotics, which form a covalent adduct with an active site serine. The enzymes responsible for 3-3 crosslinking are L,D-transpeptidases (LDTs), which are non-essential, use a catalytic cysteine rather than a catalytic serine, and are not inhibited by β-lactams, with the important exception of some penems and carbapenems^12–14^. Another critical difference is that PBPs require a pentapeptide as acyl donor for transpeptidation, while LDTs require a tetrapeptide^11,12^.

In 2011, Peltier et al. discovered that ∼75% of the crosslinks in *C. difficile* are 3-3 crosslinks, raising the intriguing possibility that 3-3 crosslinking might be essential for viability in this pathogen^15^. All known LDTs contain a YkuD catalytic domain. The *C. difficile* genome encodes three YkuD-domain LDTs, which have been characterized to various degrees both *in vivo* and *in vitro*^15–18^. Surprisingly, mutants lacking various combinations of the three LDTs are viable and synthesize PG with normal^18^ or slightly reduced 3-3 crosslinking^15^. These intriguing reports leave open two major questions: Does *C. difficile* require 3-3 crosslinked PG for viability? What enzyme(s) catalyze 3-3 crosslinking in the absence of the known YkuD-family LDTs?

Here we show 3-3 crosslinks are essential for viability in *C. difficile* and that the “missing” LDTs are two previously uncharacterized VanW domain proteins. The function of VanW domains was until now unknown. They are found primarily in Gram-positive bacteria and presumed to be involved in vancomycin resistance because of their occurrence in some atypical *Enterococcus* vancomycin resistance gene clusters. VanW domains are structurally and evolutionarily unrelated to YkuD domains. Nevertheless, in a remarkable example of convergent evolution, the presence of a conserved and essential cysteine suggests transpeptidation by VanW domains involves a thioacyl enzyme-substrate intermediate, as previously shown for YkuD domains. Our findings provide a mechanistic rationale for the occurrence of VanW domain proteins in vancomycin-resistance gene clusters and suggest that LDTs are promising targets for narrow-spectrum antibiotics against *C. difficile*.

## RESULTS

### Loss of the known Ldts has no effect on the level of 3-3 crosslinks or cell viability

All known LDTs contain a catalytic YkuD domain, which is named after a *Bacillus subtilis* protein from which one of the first crystal structures was reported^12,19^. *C. difficile* contains three YkuD-type LDTs (Fig. 2a), and these have been characterized to various degrees *in vivo* and *in vitro*^15,17,18^. To address their contributions to PG biogenesis, we used CRISPR mutagenesis to delete the three *ldt* genes alone and in combination. We had no difficulty constructing a triple deletion strain, referred to here for simplicity as *Δldt1-3* even though the genes are not in an operon. We verified the triple mutant by PCR across each *ldt* deletion, Western blotting, and whole genome sequencing, which confirmed the three *ldt* deletions are the only mutations in the strain (Supplemental Fig. 1). To our surprise, the *Δldt1-3* mutant was not only viable but completely healthy as judged by growth rate, morphology, sensitivity to a collection of cell wall-targeting antibiotics, and muropeptide analysis, which revealed no decrease in 3-3 PG crosslinks as compared to wild-type (Fig. 1b-e; Supplemental Fig. 2a,b; Tables 1 and 2). Overall, our findings are consistent with the recent characterization of an independently constructed *C. difficile* triple *ldt* deletion mutant^18^ and imply that *C. difficile* must have one or more novel LDT(s).

**Fig. 2.**
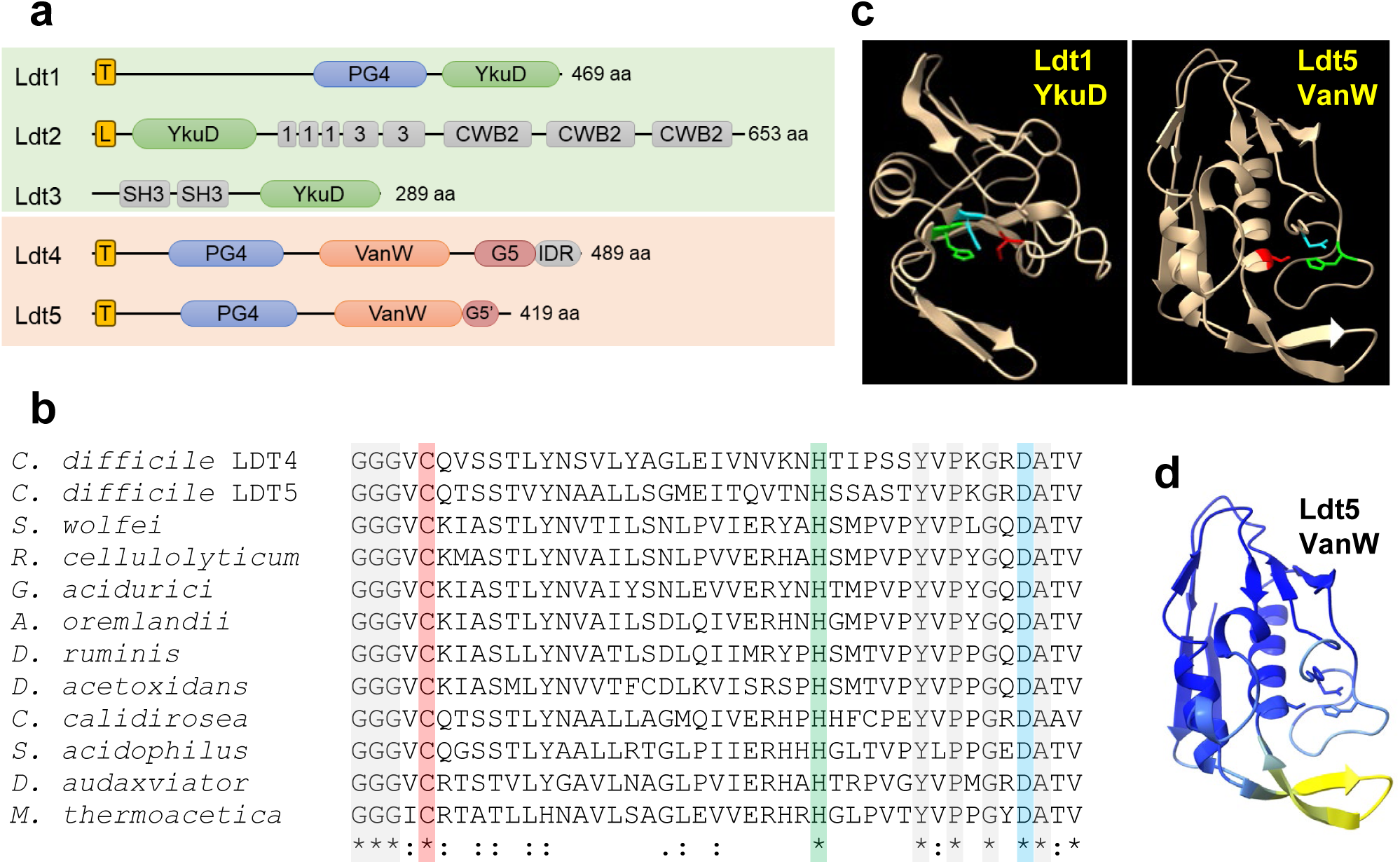
Predicted structures of *C. difficile* Ldts. (**a**) Domain architecture. T, transmembrane helix; L, signal peptidase 2 signal sequence; PG4, PG binding domain 4; YkuD, L,D-transpeptidase catalytic domain; 1 and 3, choline binding domains; CWB2, cell wall binding domain 2; SH3, Bacterial SH3 domain; VanW, L,D-transpeptidase catalytic domain; G5 and G5’, complete and partial G5 domains; IDR, intrinsically disordered region. (**b**) Amino acid sequence alignment of the active site region from 10 VanW domains with the proposed catalytic triad highlighted with red, green and blue. Gray highlight and asterisks denote strict amino acid identity, colons and periods indicate other conserved positions. Sequences shown are from *C. diffficile*, *Desulforudis audaxviator, Moorella thermoacetica, Sulfobacillus acidophilus, Ruminiclostridium cellulolyticum, Gottschalkia acidurici, Alkaliphilus oremlandii, Syntrophomonas wolfei, Desulforamulus ruminis, Desulfofarcimen acetoxidans*, and *Chthonomonas calidirosea*. See Supplementary Fig. 1 for an alignment of the entire VanW domains. (**c**) Alphafold2 models of the YkuD domain from *C. difficile* Ldt1 and the VanW domain from Ldt5, with the catalytic triads in color: Cys (red), His (green), Asp (cyan). (**d**) Confidence of the Ldt5 VanW domain model based on predicted local distance difference test. Dark blue >90 (highly accurate), light blue 89-70 (modeled well), yellow 69-50 (low confidence, caution).

**Table 1.**
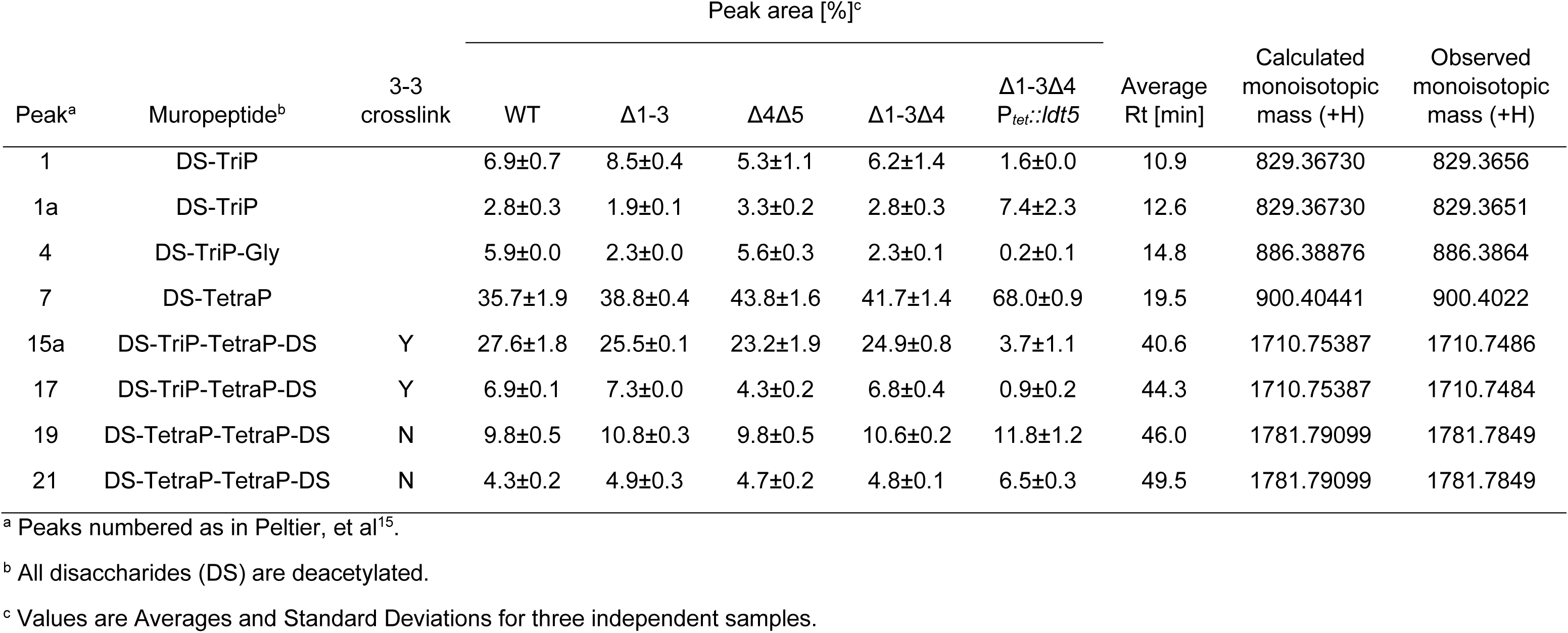
Muropeptide quantitation.

**Table 2.**
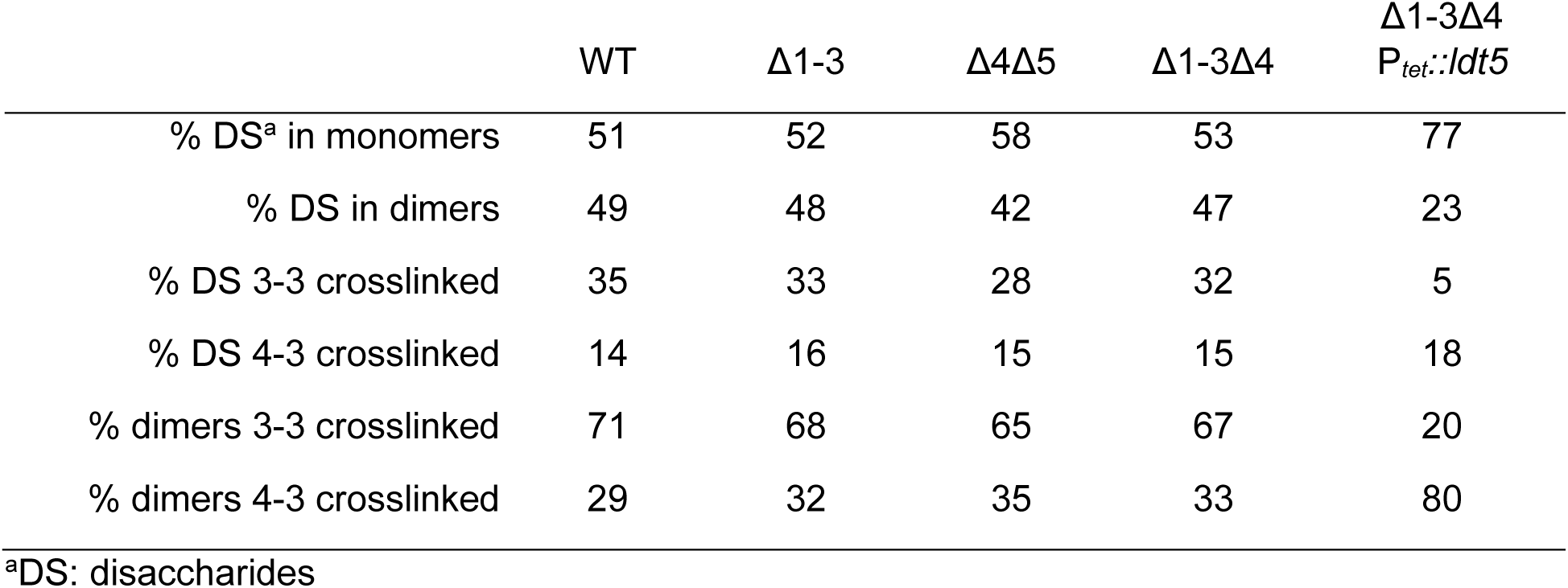
PG crosslinking.

| | WT | $\Delta 1-3$ | $\Delta 4\Delta 5$ | $\Delta 1-3\Delta 4$ | $\Delta 1-3\Delta 4$<br>$P_{tet.::ldt5}$ |
| --- | --- | --- | --- | --- | --- |
| % DS<sup>a</sup> in monomers | 51 | 52 | 58 | 53 | 77 |
| % DS in dimers | 49 | 48 | 42 | 47 | 23 |
| % DS 3-3 crosslinked | 35 | 33 | 28 | 32 | 5 |
| % DS 4-3 crosslinked | 14 | 16 | 15 | 15 | 18 |
| % dimers 3-3 crosslinked | 71 | 68 | 65 | 67 | 20 |
| % dimers 4-3 crosslinked | 29 | 32 | 35 | 33 | 80 |

<sup>a</sup>DS: disaccharides

### Bioinformatic identification of VanW domain proteins as potential LDTs

We reasoned that the missing LDT(s) might be upregulated to compensate for the absence of the three YkuD-type LDTs, so we used RNA sequencing to compare the gene expression profile of wild-type to Δ*ldt1-3* mutant. The only noteworthy differences were the absence of transcripts for the three *ldt* genes deleted in the mutant (Supplemental Fig. 2c; Supplemental Table 1). Although this experiment failed in its original goal of finding the missing LDT(s), the absence of a cell envelope stress response underscores the basic health of the Δ*ldt1-3* mutant.

We then searched the *C. difficile* R20291 genome in BioCyc^20^ for proteins with the noncatalytic domains annotated in the various *C. difficile* LDTs (Fig. 2a). Hits to the choline-binding, cell wall-binding and bacterial SH3 domains in Ldt2 and Ldt3 returned proteins that were either unlikely candidates for an LDT (e.g., PG hydrolases and the major toxins TcdA and TcdB) or too numerous to test (e.g., 27 proteins with the cell wall-binding 2 domain). In contrast, a search with the PG_binding_4 domain (PG4; PF12229) found in Ldt1 only returned two uncharacterized proteins, CDR20291_1285 and CDR20291_2055, which we have named Ldt4 and Ldt5 based on results shown below. Interestingly, all three *C. difficile* PG4 domain proteins (i.e., Ldt1, Ldt4 and Ldt5) are upregulated in a mutant lacking the PrkC serine/threonine kinase involved in cell envelope homeostasis^21^, suggesting a shared function.

Both Ldt4 and Ldt5 are predicted membrane proteins with large extracellular domains that include a PG4 domain and a VanW domain (PF04294) (Fig 2a). In Ldt4 the VanW domain is followed by a G5 domain (PF07501) and an intrinsically disordered region (IDR). Ldt5 appears to have the first half of a G5 domain (G5’). PG4 domains are often found in YkuD-type LDTs and have been proposed to bind PG^22^, but PG-binding has not been demonstrated. G5 domains are named for conserved glycines. They are found in many extracellular proteins from Gram-positive organisms^22^, bind zinc and heparin^23,24^, and have been modeled into the structure of PG^25^, but there is no experimental evidence for PG-binding. C-terminal IDRs in PBPs target these enzymes to gaps in the sacculus to make repairs^26^.

No function has been proposed for VanW domains, which were first recognized in atypical *Enterococcus* vancomycin resistance gene clusters^27,28^. Close inspection of VanW domain sequence alignments revealed a conserved cysteine, histidine and aspartate that are positioned to form a catalytic triad when mapped on to the high-confidence AlphaFold2 (AF2)^29^ models of the VanW domains from Ldt4 and Ldt5 (Fig. 2b-d; Supplemental Fig. 3,4). Similar triads are found in YkuD-type LDTs (modeled for *C. difficile* Ldt1 in Fig. 2c), which catalyze a two-step transpeptidation reaction via a covalent thioacyl-enzyme intermediate^12,30,31^. Despite this similarity, VanW and YkuD domains have different folds and cannot be superimposed. Moreover, searches with DALI and Foldseek indicate the VanW domain has no significant similarity to any known structures^32–34^. Thus, the VanW domain appears to represent a novel fold with a catalytic triad as found in YkuD-family LDTs.

### VanW domain proteins Ldt4 and Ldt5 are L,D-transpeptidases *in vitro*

Transpeptidase activity can be assayed by monitoring incorporation of fluorescent substrate analogs into isolated PG sacculi^35^. We purified the soluble extracellular domains of Ldt4, Ldt5 and as a control the YkuD domain protein Ldt1 (Ldt4^28–489^, Ldt5^38–419^, Ldt1^39–469^) (Supplemental Fig. 5a). The enzymes were tested using TetraRh, a fluorescent analog of an authentic LDT acyl donor substrate^36^. TetraRh consists of a Rhodamine dye attached to the N-terminus of a tetrapeptide based on *E. faecium* PG with the sequence: D-Ala-iso-DGln-L-Lys(Ac)-D-Ala (Fig. 3a). Note that the Lysine is acetylated to prevent it from acting as a transpeptidation acceptor. To test for LDT activity, enzyme and TetraRh were incubated with *Bacillus subtilis* PG sacculi that had been immobilized on a glass slide. After 1hr, the sacculi were washed and imaged by phase contrast and fluorescence microscopy. The results are compiled in Fig. 3b. All three LDTs incorporated TetraRh into sacculi, although fluorescence was about 4-fold higher with Ldt1 than Ldt4 or Ldt5 (Fig. 3c). In contrast, none of the LDTs incorporated label when D-Ala^4^ in TetraRh was replaced with L-Ala, nor did they incorporate a pentapeptide analog (PentaRh) that is a substrate for PBPs but not LDTs^36^. As expected, changing the active site cysteine to alanine in Ldt4 and Ldt5 abrogated activity with TetraRh. Circular dichroism spectra of the Ldt4^C286A^ and Ldt5^C298A^ mutant proteins were indistinguishable from WT, indicating they folded properly (Supplemental Fig. 5b). Finally, meropenem inhibited the activity of Ldt1, Ldt4 and Ldt5 (Fig. 3d). Inhibition of Ldt1 was expected as meropenem is known to acylate this enzyme^17^. Inhibition of Ldt4 and Ldt5 argues VanW domains and YkuD domains have very similar active sites, which has implications for developing antibiotics effective against both LDT families.

**Fig. 3.**
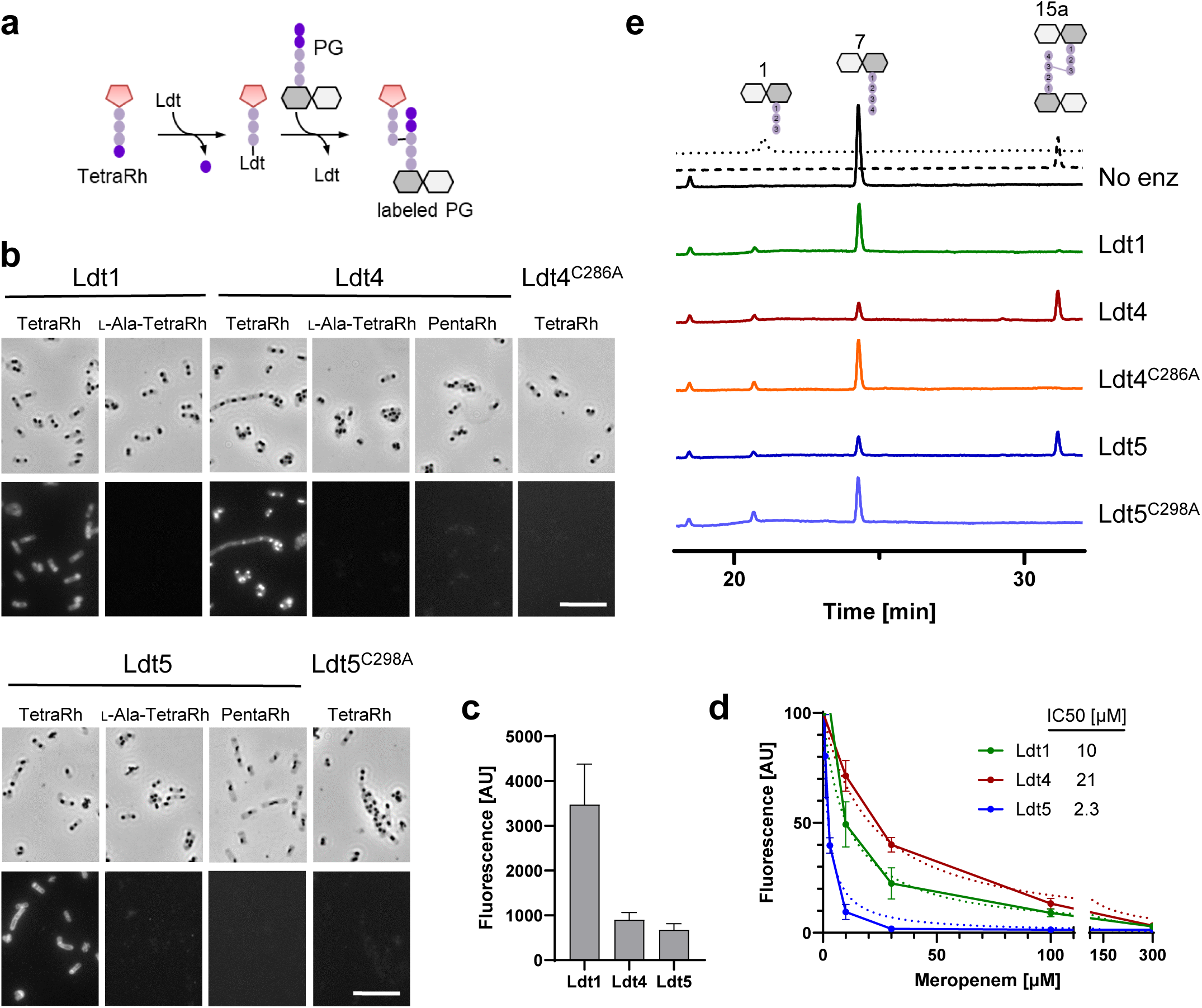
VanW domains catalyze L,D-transpeptidation *in vitro*. (**a**) Schematic diagram of incorporation of TetraRh into PG sacculi by an Ldt. Pink pentagon, Rhodamine. Colored balls, amino acids. Dark and light gray hexagons, NAM and NAG, respectively. (**b**) Phase contrast and fluorescence micrographs of immobilized PG sacculi after incubation for 1 h with 5 μM enzyme and 30 μM substrate analog as indicated. TetraRh: LDT-specific substrate analog; L-Ala-TetraRh, negative control; PentaRh, PBP-specific substrate. Size bar, 10 μm. Micrographs are representative of at least two experiments. (**c**) Quantification of TetraRh incorporation into sacculi graphed as the mean ± s.d. of the fluorescence intensity from 10 sacculi. (**d**) Inhibition of LDT activity by meropenem graphed as the mean and s.d. of data pooled from four experiments. IC50 is the concentration of meropenem needed to reduce LDT activity by half. (**e**) HPLC analysis of muropeptides after 1 h incubation of the indicated enzymes with DS-TetraP substrate. Structures above the chromatograms are numbered according to Peltier et al.^15^. Calibration traces for 1 and 15a are shown by the dotted and dashed lines, respectively. Chromatograms are representative of 3 experiments.

We next tested Ldt4 and Ldt5 for L,D-transpeptidase activity using disaccharide-tetrapeptide (DS-TetraP) isolated by HPLC after mutanolysin digestion and borohydride reduction of purified *C. difficile* PG sacculi. DS-TetraP has the structure NAG-NAM(red)-L-Ala-iso-D-Glu-*meso*-DAP-D-Ala, with the NAM moiety in the muramitol form due to the reduction step. Reaction mixtures containing enzyme and DS-TetraP were incubated for 2 h at 37°C, then analyzed by HPLC. Ldt4 and Ldt5 converted DS-TetraP to a product with a retention time of 31 min (Fig. 3e), which was determined by elution time and mass spectrometry to be disaccharide-tripeptide 3-3 crosslinked to a disaccharide-tetrapeptide (DS-TriP-TetraP-DS) (Supplemental Fig. 6). As expected, the catalytic mutant derivatives Ldt4^C286A^ and Ldt5^C298A^ were unable to produce a crosslinked product, although both generated some DS-TriP, indicating they retain carboxypeptidase activity (Fig. 3e). We interpret these results to mean that the catalytic cysteine is required for transpeptidation, presumably because of its role as an acyl carrier, but other features of the enzyme that promote catalysis such as transition state stabilization are sufficient for removal of the terminal D-Ala^4^. Of note, but in agreement with a previous study^17^, Ldt1 did not crosslink DS-TetraP in our assay, although Ldt1 has been shown to generate a small amount of crosslinked product by using more enzyme and longer incubations^18^. The fact that Ldt1 has little or no ability to crosslink DS-TetraP yet outperforms Ldt4 and Ldt5 for incorporation of TetraRh into intact sacculi suggests it requires a substrate larger than a DS-TetraP as acceptor in the transpeptidation reaction.

### LDTs and 3-3 crosslinks are essential for viability in *C. difficile*

Having determined that the VanW domain proteins are *bona fide* LDTs, we addressed their contribution to 3-3 crosslinking *in vivo*. To this end, we used CRISPR mutagenesis to create the *Δldt4*, *Δldt5*, and *Δldt4Δldt5* strains. All of these mutants were viable and exhibited typical rod morphology (Fig. 4a; Supplemental Fig. 7a). In addition, muropeptide analysis of the *Δldt4Δldt5* strain revealed normal levels of 3-3 crosslinking (Fig. 5a; Tables 1 and 2). We then deleted either *ldt4* or *ldt5* in the *Δldt1-3* background. Once again the mutants were viable with no change in morphology (Fig. 4a; Supplemental Fig. 7a), and the Δ*ldt1-3Δ4* strain retained normal levels of 3-3 crosslinking (Fig. 5a; Tables 1 and 2). We also tested if sporulation was affected in some of the mutants and observed a modest increase in some cases (Supplemental Fig. 7b). However, multiple attempts to delete all five *ldt*s were unsuccessful, suggesting a synthetic lethal phenotype. This was confirmed using CRISPRi to knock down expression of *ldt4* in a *Δldt1-3Δ5* mutant or *ldt5* in a *Δldt1-3Δ4* mutant. In both cases knockdown of the last remaining LDT caused a 2-3 log drop in viability (Supplemental Fig. 8)

**Fig. 4.**
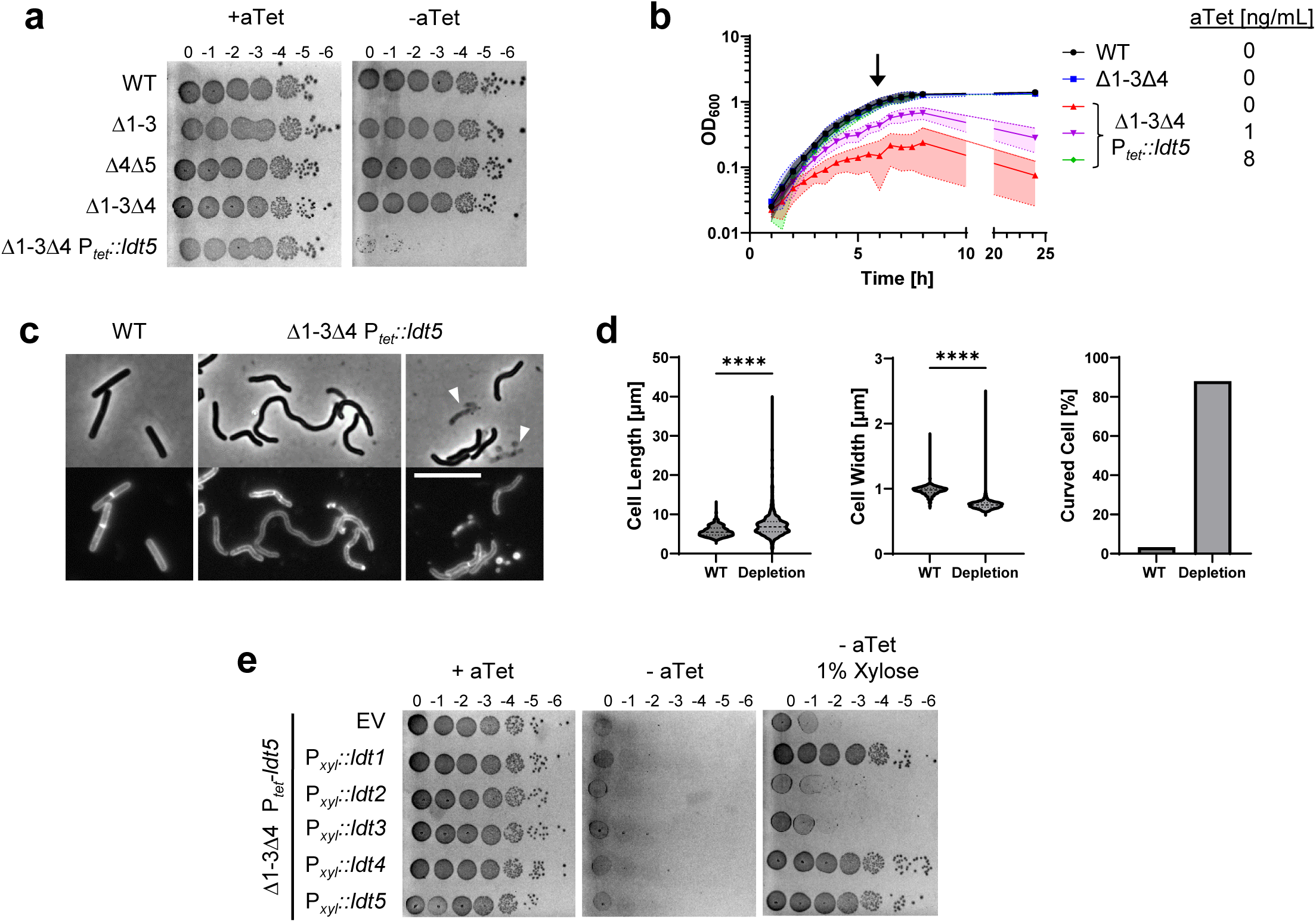
LDTs are essential in *C. difficile*. (**a**) Viability assay. Ten-fold serial dilutions of the indicated strains were spotted onto TY plates with or without 25 ng/mL aTet. Plates were photographed after incubation for 18 h. Images are representative of at least three experiments. (**b**) Growth curves. Data are graphed as the mean ± s.d. of 4 biological replicates from different days. (**c**) Cell morphology. Cells grown for 6 h in TY without aTet (arrow in B) were stained with the membrane dye FM4-64 and photographed under phase contrast and fluorescence microscopy. Arrowheads indicate lysed cells. Size bar, 10 μm. Images representative of at least three experiments. (**d**) Quantification of length, width and shape based on 781 cells of WT and 1196 cells of the depletion strain pooled from three biological replicates. Cells with a sinuosity score ≥1.03 were considered curved. ****, p < 0.0001, unpaired *t*-test. (**e**) Complementation assay. Tenfold serial dilutions of the LDT depletion strain harboring the indicated expression plasmids were spotted onto TY with or without 25 ng/mL aTet and 1% xylose. Plates were photographed after incubation for 18 h. Images are representative of three biological replicates. Strains shown in A-D: WT, R20291; Δ1-3, KB124; Δ4Δ5, KB529; Δ1-3Δ4, KB474;and Δ1-3Δ4 P*_tet_*::*ldt5*, KB547 (called “depletion” in panel **d**). Strains shown in panel **e**: empty vector (EV), KB548, P*_xyl_::ldt1,* KB549; P*_xyl_::ldt2,* KB550; P*_xyl_::ldt3,* KB551; P*_xyl_::ldt4,* KB552; and P*_xyl_::ldt5,* KB553.

**Fig. 5.**
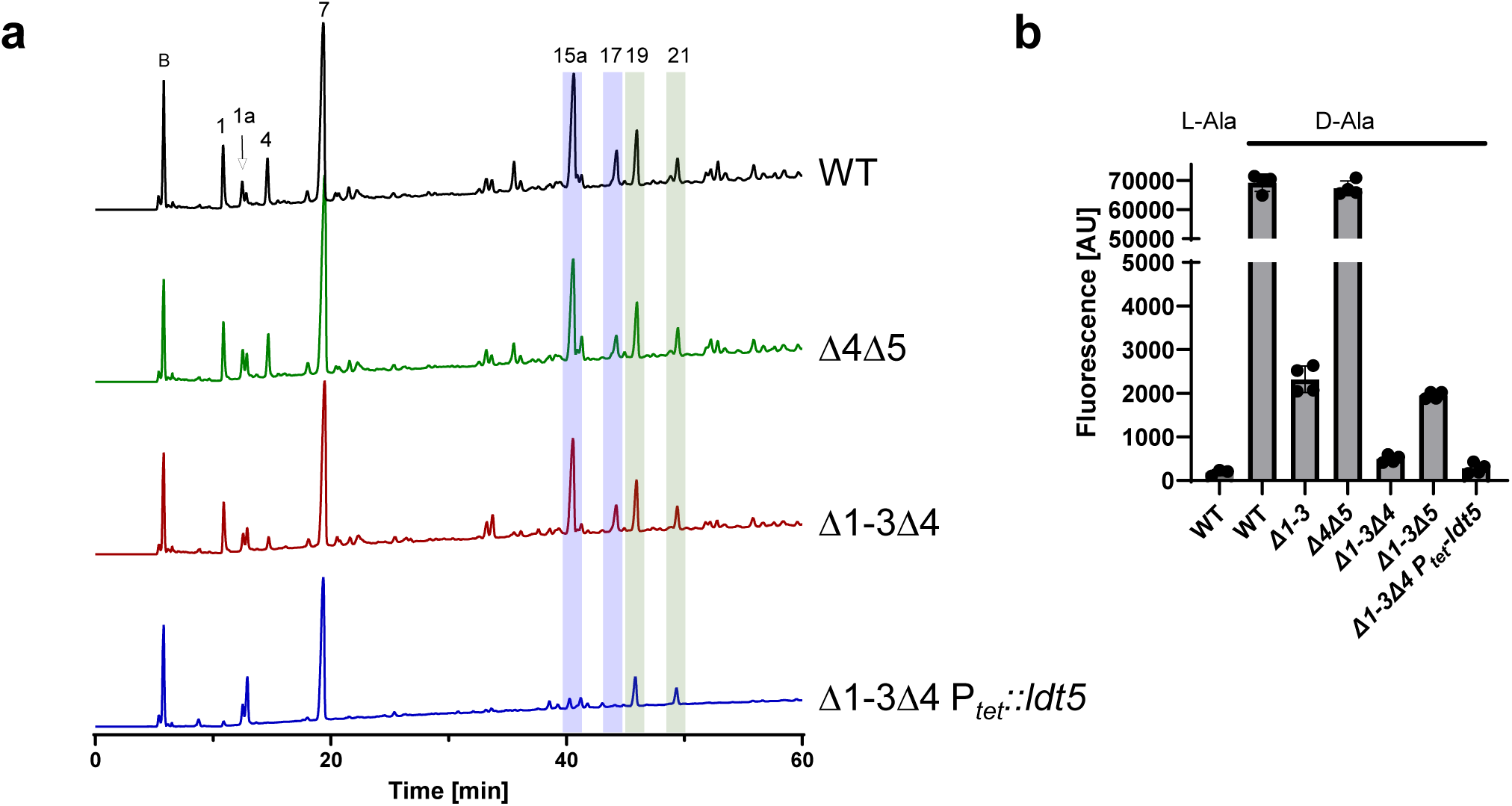
YkuD and VanW LDTs create 3-3 crosslinks *in vivo*. (**a**) HPLC quantitation of muropeptides from the indicated strains. Blue and green highlights indicate the major 3-3 and 4-3 crosslinked muropeptides, respectively. Peak numbers based on Peltier et al.^15^ (**b**) Flow cytometry analysis of cells grown for 1 h in the presence of TetraRh. L-Ala is a control and refers to a non-physiological TetraRh analog with L-alanine rather than D-alanine in position 4. Strains: WT, R20291; Δ4Δ5, KB529; Δ1-3Δ4, KB474; Δ1-3Δ4 P*_tet_::ldt5,* KB547; Δ1-3, KB124; and Δ1-3Δ5, KB502.

We then replaced the native promoter for *ldt5* with P*_tet_* in the *Δldt1-3Δ4* background, rendering expression of the last remaining LDT dependent on the inducer anhydrotetracycline (aTet). Spot titer assays on TY media with and without aTet revealed Ldt5 was required for viability (Fig. 4a). This result was confirmed in liquid media, where subculturing into TY lacking aTet resulted in slower growth and eventually a drop in OD_600_ indicative of lysis (Fig 4b). Microscopy revealed cells depleted of the last remaining LDT became longer, thinner and curvy in comparison to WT. Cell ghosts indicative of lysis were also seen (Fig. 4c,d). Staining with the membrane dye FM4-64 revealed relatively few division septa in the population depleted of LDTs (Fig. 4c). These phenotypic defects implicate LDTs in elongation and cell division.

To determine whether the viability loss is associated with the loss of 3-3 crosslinks, we analyzed muropeptides from WT, the *Δldt4*Δ*ldt5* double mutant, the *Δldt1-3Δ4* quadruple mutant, and the *Δldt1-3Δ4* P*_tet_*::*ldt5* depletion strain. Cultures were grown in TY, which was supplemented with a small amount of aTet (0.25 ng/mL) for the depletion strain so it could reach high enough OD_600_ to obtain sufficient sacculi for muropeptide analysis. Muropeptides were identified by mass spectrometry and named according to Peltier et al. to facilitate comparisons (Fig. 5a)^15^. There was a drastic reduction in muropeptides 1, 4, 15a and 17 in the depletion strain. All of these changes are attributable to loss of LDT activity. Most importantly, peaks 15a and 17 are two 3-3 crosslinked DS-TriP-TetraP-DS species that separate during HPLC for unknown reasons. The area under the curve for muropeptides 15a + 17 decreased from 34.5% in WT to 4.6% in the depletion strain, an ∼85% decrease (Tables 1 and 2). Peaks 1 and 4 are DS-TriP and DS-TriP-Gly, which are created by LDT-catalyzed carboxypeptidase and exchange reactions, respectively. A minor DS-TriP species eluting as peak 1a appears to be increased in the depletion stain; we cannot at present explain that change. Interestingly, loss of 3-3 crosslinking increased primarily uncrosslinked muropeptides (Peaks 1a and 7) rather than 4-3 crosslinking muropeptides (Peaks 19 and 21). Thus, LDTs and PBPs are not in competition, which can be explained by their different acyl donor requirements, and *C. difficile* is unable to compensate for defects in 3-3 crosslinking by making more 4-3 crosslinks instead.

We further characterized LDTs in vivo using TetraRh (Fig. 5b). Flow cytometry of wild type *C. difficile* cells grown for 1 h in the presence of 30 μM TetraRh revealed a ∼350-fold increase in fluorescence as compared to background determined using the non-physiological L-Ala-TetraRh variant. Label incorporation decreased ∼30-fold in the Δ*ldt1-3* strain but was unaffected in the Δ*ldt4*Δ*ldt5* strain, indicating TetraRh is a much better substrate for *C. difficile’s* YkuD-type LDTs than for its VanW-type LDTs, as was seen *in vitro* (Fig 3b,c). Deletion of *ldt4* in the Δ*ldt1-3* background further reduced TetraRh incorporation, but deletion of *ldt5* had little effect. Finally, labeling with TetraRh dropped to near background when the *Δldt1-3Δ4* P*_tet_*::*ldt5* depletion strain was grown without aTet (Fig. 5b). Overall, experiments with TetraRh confirm the absence of LDT activity in the *Δldt1-3Δ4* P*_tet_*::*ldt5* depletion strain.

### Ldt1, Ldt4 or Ldt5 is sufficient for viability

The above results demonstrate *C. difficile* must express at least one *ldt* for 3-3 crosslinking and viability. But which one(s)? To address this question, we cloned each *ldt* into a plasmid with a xylose-inducible promoter, P*_xyl_*^37^. The resulting *ldt* expression plasmids were conjugated into the *Δldt1-4* P*_tet_*::*ldt5* depletion strain, and viability was determined by a spot titer assay on TY with 1% xylose but no aTet. We found that Ldt1, Ldt4, or Ldt5 were each sufficient for viability, but Ldt2 or Ldt3 were not (Fig. 4e), even though they were produced at physiological levels or higher as determined by Western blotting (Supplemental Fig. 1b-d). Moreover, CRISPRi knockdown of *ldt1* in a *Δldt4-5* mutant resulted in a loss of viability, indicating that Ldt2 and Ldt3 are not sufficient for normal growth even when present simultaneously (Supplemental Fig. 8).

### VanW domain containing proteins are common in Gram-positive bacteria

The Pfam database (v31, November 2023) lists 15,131 VanW domain proteins from 6920 bacterial species^38^. That makes VanW domains almost 10-fold less common than YkuD domains, for which Pfam lists about ∼131,000 examples in ∼30,000 bacterial species. About half of the VanW domain proteins have one or more PG4 domains, as seen in Ldt4 and Ldt5 of *C. difficile*. Using AnnoTree^39^ to map the Pfam VanW domains onto a bacterial phylogenetic tree revealed that ∼70% of Bacillota (formerly called Firmicutes) and ∼40% of Actinomycetota encode at least one VanW domain protein (Supplemental Fig. 9). Indeed, these two phyla account for ∼65% of all sequenced VanW homologs. VanW domains are also relatively common in the Chloroflexota and Patescibacteria. In contrast, only ∼10% of Cyanobacteria, 6% of Bacteriodiota, and 1% of Pseudomonadota genomes encode a VanW domain protein.

To ask whether LDT activity is a common property of VanW domain proteins, we tested seven (Fig. 6a) for the ability to incorporate TetraRh when expressed in *C. difficile*. As a positive control for this experiment, we used *C. difficile* Ldt5. All eight genes were expressed from a P*_xyl_* plasmid in the *C. difficile* Δ*ldt1-3*Δ*ldt4* mutant, where the background level of TetraRh incorporation is quite low (Fig. 5b). Three of the seven foreign VanW domain proteins supported incorporation of TetraRh into *C. difficile*, indicating they are indeed LDTs (Fig. 6b). Negative results in this assay are not readily interpreted because, for example, we do not know if the apparently inactive proteins were produced.

**Fig. 6:**
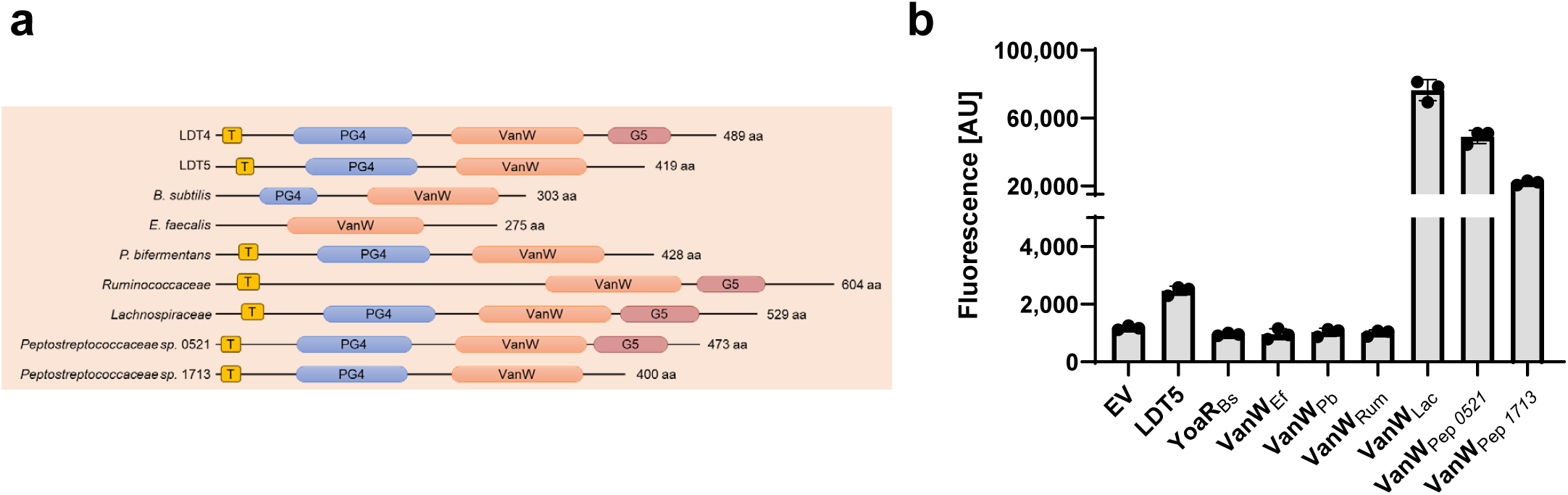
Survey of VanW domain proteins from various Bacillota for LDT activity. (**a**) Domain structures labeled as in Fig. 1a. (**b**) Flow cytometry analysis of cells grown for 1 h in the presence of TetraRh graphed as the mean ± s.d. from 3 trials. The strains are derivatives of KB474 [Δ*ldt1-3*Δ*ldt4*] harboring P*_xyl_* expression vectors: EV, KB633; *C. difficile* Ldt5, KB634; *B. subtilis* YoaR, KB635; *Enterococcus faecalis* VanW, KB636; *Paraclostridium bifermentas* VanW, KB637; *Ruminococcaceae sp.* VanW, KB638; *Lachnospiraceae sp.* VanW, KB639; *Peptostreptococcus* sp. VanW 0521, KB640; and *Peptostreptococcus* sp. VanW 1713, KB 641.

## DISCUSSION

Most well-studied bacteria rely primarily on PBPs that make 4-3 crosslinks to construct an osmotically stable PG wall. *C. difficile*, in contrast, relies primarily on 3-3 crosslinks created by LDTs^15^. In this paper we have demonstrated that 3-3 crosslinking and LDTs are essential for viability in *C. difficile*, making it the first and so far the only bacterium in which 3-3 crosslinks and LDTs are known to be essential. We also report the discovery of a new family of LDTs that employ a VanW catalytic domain, which has no sequence or structural similarity to the YkuD catalytic domain found in all previously known LDTs^12^. Indeed, VanW domains appear to represent a novel fold, as searches to detect related structures using Foldseek^32^ or remote homologs using HHsearch^40^ did not return any statistically significant matches. Nevertheless, we infer that VanW and YkuD domains catalyze 3-3 crosslinking by a similar two-step catalytic mechanism based on the fact that both domains have a conserved cysteine that is required for transpeptidation. In YkuD domains this cysteine is the attacking nucleophile that forms a covalent thioacyl intermediate with the donor peptide substrate^13,30,41^.

VanW domains are named for their presence in atypical *Enterococcus* vancomycin resistance gene clusters^27,28^. There is no experimental evidence for a role in vancomycin resistance, nor has any biochemical function been proposed. Our discovery that VanW domains catalyze 3-3 crosslinking suggests a mechanism by which they could contribute to vancomycin resistance. Vancomycin inhibits PG synthesis by binding to the terminal D-alanyl-D-alanine of the pentapeptide in PG precursors. Known resistance mechanisms involve modifying the stem peptide to prevent vancomycin binding by changing the terminal D-alanine to D-serine or D-lactate^42,43^, or by converting pentapeptides to tetrapeptides that are subsequently crosslinked by LDTs^44^. But the latter resistance mechanism comes at a cost because it renders the PBPs inoperative. Curiously, *C. difficile* is vancomycin sensitive despite its heavy reliance on LDTs for PG crosslinking^15,45^. Further work will be needed to understand this conundrum, but it could have to do with the fact that *C. difficile* has two PBPs that are essential for vegetative growth. Alternatively, or in addition, vancomycin might block conversion of pentapeptides to tetrapeptides by extracellular carboxypeptidases and thus starve LDTs of substrate. Similarly, dual inhibition of synthetic PBPs and carboxypeptidases might explain why *C. difficile* is sensitive to β-lactams like ampicillin that do not inhibit LDTs directly^46^.

In considering the potential advantages of LDTs over PBPs in PG biogenesis, an important distinction is that only LDTs can repair broken crosslinks in the absence of *de novo* PG synthesis^47^. In particular, endopeptidase cleavage of a 4-3 crosslink generates tetra- and tripeptides that can be stitched back together as a 3-3 crosslink by an LDT but not a PBP. The repair function of LDTs is important for maintaining PG integrity in *Mycobacterium smegmatis* and presumably other bacteria that exhibit polar growth and high levels of 3-3 crosslinking^47^. However, we hypothesize that *C. difficile* employs LDTs as the major source of initial crosslinking during elongation and perhaps division as well. Using the fluorescent D-amino acid HADA to visualize sites of PG synthesis in growing *C. difficile* cells revealed uniform incorporation throughout the sidewall, arguing against polar growth^48^. Moreover, the morphological defects we observed upon LDT depletion—longer, thinner cells with few septa—are more suggestive of a PG synthesis defect than a repair defect, which should have manifested as bloating and bulges, as reported in *M. smegmatis* ^47^.

The unique essentiality of 3-3 crosslinks in *C. difficile* suggests LDTs should be explored as targets for antibiotics that kill *C. difficile* without disrupting the normal intestinal microbiota needed to keep *C. difficile* in check. Previous efforts to develop antibiotics that inhibit LDTs have focused mainly on *Mycobacterium tuberculosis* Ldt_Mt2_, which is required for virulence but not for viability *per se*^49,50^. These efforts have mostly been directed at improving the efficacy of penems and carbapenems^41,51,52^. However, penems and carbapenems also inactivate PBPs. This may be a plus for treating tuberculosis but compromises the selectivity that makes LDTs attractive therapeutic targets in *C. difficile*. Nevertheless, our finding that meropenem inhibits *C. difficile’s* YkuD and VanW domain LDTs argues that it is possible to develop drugs that target both classes of LDTs despite their profoundly different structures.

## Supporting information

Supp Table 1_RNA-seq

## ACKNOWLEDGMENTS

The following reagent was obtained through the ATCC: *Paraclostridium bifermentans* ATCC 638. The following reagents were obtained through BEI Resources, NIAID, NIH as part of the Human Microbiome Project: *Eubacterium* sp., Strain AS15, HM-766; *Ruminococcaceae* sp., Strain D16, HM-79; *Lachnospiraceae* sp., Strain 5_1_57FAA, HM-157.

This work was supported by the National Institutes of Health (R01AI155492 to C.D.E. and D.S.W., R01GM138630 to D.L.P., and R35GM124893-06 to M.M.P.) The LCMS work was funded by GlycoMIP, a National Science Foundation Materials Innovation Platform funded through Cooperative Agreement DMR-1933525. Cell fluorescence was quantitated at the Flow Cytometry Facility, a core research facility at the University of Iowa funded through user fees and the generous financial support of the Carver College of Medicine, Holden Comprehensive Cancer Center, and Iowa City Veteran’s Administration Medical Center. RNA integrity was characterized by the Genomics Division of the Iowa Institute of Human Genetics, which is supported, in part, by the University of Iowa Carver College of Medicine. We thank Johann Peltier for sharing information prior to publication, Atsushi Yahashiri for sharing *B. subtilis* PG sacculi and members of the Ellermeier and Weiss laboratories for helpful discussions.

## AUTHOR CONTRIBUTIONS

K.W.B., U.M. and C.D.E. constructed plasmids. K.W.B. constructed and characterized *C. difficile* strains, performed RNA-seq, and performed in vivo assays involving TetraRh and derivatives. C.D.E. performed bioinformatics. U.M. purified LDTs and assayed LDT activity with TetraRh in vitro. K.L.O and A.J.A. synthesized TetraRh and derivatives. M.M.P. supervised K.L.O. and A.J.A. and provided guidance on use of TetraRh and derivatives. R.F.H. and D.L.P. performed muropeptide analyses. D.L.P. assayed crosslinking of DS-TetraP. D.S.W and C.D.E. provided overall supervision of the project. The manuscript was drafted by K.W.B, U.M., D.S.W. and C.D.E. All other authors reviewed and edited the manuscript.

## MATERIALS AND METHODS

### Strains, media, and growth conditions

Bacterial strains are listed in Table 3 and Supplemental Table 2. *C. difficile* strains used in this study were all derived from R20291^53^. *C. difficile* was grown in tryptone-yeast (TY) media, supplemented as needed with thiamphenicol at 10 μg/mL (Thi_10_), kanamycin at 50 μg/mL, or cefoxitin at 8 μg/mL. Anhydrous tetracycline (aTet) was used to induce genes under P*_tet_* control (Fluka). TY media consisted of 3% tryptone, 2% yeast extract, and 2% agar (for solid media). For conjugation plates brain heart infusion (BHI, Bacto) solid media was used. BHI media consisted of 3.7% BHI and 2% agar. *C. difficile* strains were grown at 37°C in an anaerobic chamber (Coy Laboratory Products) in an atmosphere of about 2% H_2_, 5% CO_2_, and 93% N_2_. Growth was monitored at OD_600_ with a WPA Biowave CO8000 Cell Density Meter.

**Table 3.**
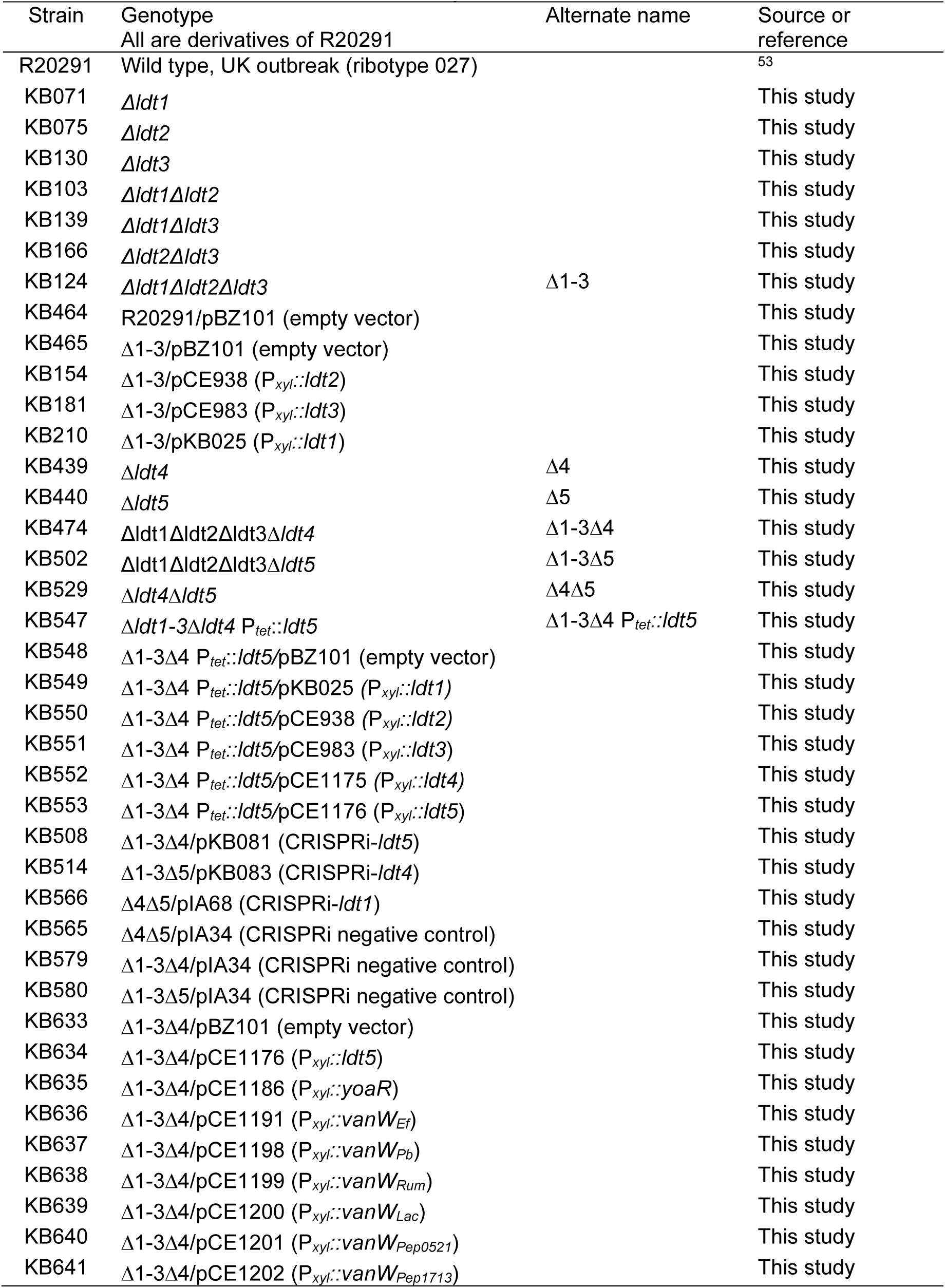
*C. difficile* strains used in this study.

*Escherichia coli* and *Bacillus subtilis* strains were grown in LB media at 37°C with chloramphenicol at 10 μg/mL or ampicillin at 100 μg/mL as necessary. LB media contained 1% tryptone, 0.5% yeast extract, 0.5% NaCl, and 1.5% agar (for solid media).

### Fluorescent substrate analogs

The L,D-transpeptidase specific substrate analog, Rhodamine-L-Ala-iso-D-Gln-L-Lys(Ac)- **D-Ala** (TetraRh), the negative control, Rhodamine-L-Ala-iso-D-Gln-L-Lys(Ac)-**L-Ala** (L-ala-TetraRh) and the PBP specific substrate analog, Rhodamine-L-Ala-iso-D-Gln-L-Lys(Ac)-**D-Ala-D-Ala** (PentaRh) were synthesized as described^36^ (Supplemental Fig. 10).

### Plasmid and bacterial strain construction

Plasmids are listed in Supplemental Table 3 and were constructed by isothermal assembly with reagents from New England Biolabs (Ipswich, MA). Regions that were constructed by PCR were verified by DNA sequencing. The oligonucleotide primers used in this study were synthesized by Integrated DNA Technologies (Coralville, IA) and are listed in Supplemental Table 4. All plasmids were propagated using OmniMax 2-T1R as the cloning host. Plasmids with the *E. coli* origin of transfer (RP4 *oriT traJ*) were transformed into HB101/pRK24^54,55^ and introduced into *C. difficile* by conjugation^56^. Plasmids which harbor the *oriT*_(Tn916)_ origin of transfer were passaged through *E. coli* MG1655, transformed into *B. subtilis* strain BS49 and conjugated into *C. difficile* R20291^57^. CRISPR editing plasmids were designed as previously described^58^ with a single guide RNA (sgRNA) against the target gene and homology regions to repair the double-stranded break caused by the Cas9 nuclease. Mutagenesis was induced by plating R20291 harboring the appropriate plasmid onto TY Thi_10_ with 1% xylose. The exact amount of cells to plate was determined experimentally by plating serial dilutions. Typical plating efficiency was around 10^-4^ but varied by construct. Survivor colonies were restruck once on TY with 1% xylose and subsequently maintained by serial passage on TY until the plasmid was lost as evidenced by sensitivity to thiamphenicol. Successful mutagenesis was confirmed by PCR with Q5 or Taq DNA polymerase (NEB). For this, template genomic DNA was prepared by picking one colony into 50 μL ThermoPol buffer (NEB) with 0.5 μL thermolabile Proteinase K (NEB), incubating at 37°C for 30 min and inactivating the protease at 55°C for 10 min.

### Antibiotic MIC determination

The MIC against select antibiotics was determined as described in biological duplicate on two separate days^56^. Briefly, overnight cultures were diluted 1:100 into TY, grown to OD_600_ ∼0.8 and then diluted to a calculated OD_600_ = 0.005 (∼10^6^ CFU/mL). A 50 μl aliquot of cells was added to 50 μl of TY plus antibiotic in 96 well plates. Growth was scored after ∼17 h incubation.

### Viability Plating

Viability was tested by a spot titer assay. For this, a 10-fold dilution series was prepared from overnight cultures, and 5 μL of each dilution were spotted on to the appropriate plates, which were incubated at 37°C overnight and imaged.

### Metabolic labeling of PG in live cells with TetraRh

For labeling live cells, most *C. difficile* strains were subcultured 1:100 into TY, grown to an OD_600_ of 0.2-0.3, and metabolic label was added to a final concentration of 30 μM. The LDT depletion strain was grown overnight in TY containing aTet at 4 ng/μl, washed once in TY to remove aTet, and then subcultured 1:100 into TY containing aTet at 0.25 ng/μl. TetraRh was added when the culture reached OD_600_ = 0.2. Typically about 1 mL of culture was incubated with dye. After 1 hour at 37°C, cells were washed three times with 1 mL phosphate-buffer saline (PBS: 137 mM NaCl, 3 mM KCl, 10 mM NaH_2_PO_4_, 2 mM KH_2_PO_4_, pH 7.4) resuspended in 100 μL PBS and fixed by pipetting into 24 μL fixation cocktail (4 μL 1M NaPO_4_ buffer, pH 7.4, and 20 μL 16% (wt/vol) paraformaldehyde (Alfa Aesar)). Fixed cells were incubated at room temperature for 30 min, then on ice for 30 min, and washed three times in PBS, suspended in ∼50 μL PBS, then imaged by microscopy or analyzed by flow cytometry.

### Microscopy

Cells were immobilized using thin agarose pads (1%). Phase-contrast micrographs were recorded on an Olympus BX60 microscope equipped with a 100× UPlanApo objective (numerical aperture, 1.35). Micrographs were captured with a Hamamatsu Orca Flash 4.0 V2+ complementary metal oxide semiconductor (CMOS) camera. Excitation light was generated with an X-Cite XYLIS LED light source. Membranes were stained with the lipophilic dye FM4-64 (Life Technologies) at 10 μg/mL. Cells were imaged immediately without washing. Red fluorescence was detected with the Chroma filter set 49008 (538 to 582 nm excitation filter, 587 nm dichroic mirror, and a 590 to 667 nm emission filter). Fluorescence was quantitated using the image analysis package Fiji^59^. The plug-in module MicrobeJ was used to measure cell length^60^.

### Flow cytometry

Fixed cells labeled with D-Ala-TetraRh or L-Ala-TetraRh were analyzed at the Flow Cytometry Facility at the University of Iowa using a Becton Dickinson LSR II instrument with a 561 nm laser, a 610/20-nm-band-pass filter, and a 600 LP dichroic filter as previously described^61^. The data were analyzed using BD FACSDiva software. Fluorescence was quantitated at 900 V and the mean reported from 20,000 cells.

### Peptidoglycan purification for muropeptide analysis

Peptidoglycan purification was adapted from previously published procedures^62,63^. Cells grown in 100 mL TY to an OD_600_ of 0.3 were pelleted by centrifugation. Pellets were resuspended in 2 mL cold water, dripped into 50 mL boiling 4% sodium dodecyl sulfate (SDS) and boiling was continued for 30 minutes. Pellets were washed with 60°C warm water at least three times or until there were no traces of SDS remaining^64^. Washed pellets were resuspended in 100 mM TrisHCl, pH 7.5, 20 mM MgCl_2_ and digested with 10 μg DNAse I (NEB) and 50 μg RNAse (Sigma-Aldrich) for 2 hours at 37°C. Samples were then adjusted to 10 mM CaCl_2_ and further digested with 100 μg trypsin (TPCK-treated, Worthington) at 37°C overnight. Teichoic acids were removed by resuspending the pellets in 6N hydrochloric acid (VWR, 50% v:v) and rocking for 48 hours at 4°C. Samples were washed 3 times in water, then incubated with 5 units of Antarctic phosphatase (NEB) at 37°C overnight. Finally, phosphatase was inactivated by heating to 95°C for 5 min, and samples were again washed 3 times with water, before storage at -20°C as a pellet.

### Muropeptide analysis

PG purified from 100 ml of culture was digested with 125 units of Mutanolysin (Sigma) in 12.5 mM NaPO_4_ pH 5.5 at 37°C for 16 hours. Insoluble material was removed by centrifugation at 15,000 x g for 10 minutes, and the supernatant containing muropeptides was lyophilized. Muropeptides were reduced using NaBH_4_ and separated using HPLC with detection at 206nm as previously described^65^. Muropeptide peaks were collected and further purified individually on the HPLC using a volatile buffer containing acetonitrile and trifluoroacetic acid^65^. These purified muropeptide peaks were collected, lyophilized, and used for mass spectrometry analyses and for *in vitro* analyses of LDT activity. LC-MS was performed as described^66^ by reversed phase chromatography (Waters BEH C18) using acidified water/MeOH gradients with the column eluents evaluated by ESI in the positive ion mode on both a Shimadzu LCMS 9030 (QTof) as well as a Bruker timsTOF FleX MALDI-2.

### Western blot

Rabbit polyclonal antiserum against Ldt1, Ldt2, and Ldt3 was raised against purified protein (ProSci). To analyze protein levels by Western immunoblotting, 3 mL cultures grown to an OD_600_ ∼ 0.85 were pelleted, resuspended in 300 μL 2x Laemmli buffer and sonicated (Branson Sonifier 450, microtip, output 3, two cycles of 15 pulses). After heating at 95°C for 10-15 min, 20 μL sample were electrophoresed on 10% SDS-PAGE (TGX gel, BioRad), transferred to nitrocellulose and developed using standard laboratory procedures. Primary antiserum for Ldt1 and Ldt3 was used at 1:10,000, primary antiserum for Ldt2 was used at 1:100,000. Secondary antibody (IRDye 680LT goat anti-rabbit antibody, LI-COR, Lincoln, NE) was used at 1:10,000 and blots were visualized with an Azure Biosystems Sapphire Biomolecular Imager.

### Sporulation

Effects of *ldt* mutation on the ability to sporulate were measured as previously described^67^.

### Production and purification of L,D-transpeptidases

Expression strains were grown in 1 L LB Amp^100^ Cm^10^ at 37°C to an OD_600_ of 0.5, induced with 1 mM IPTG, shifted to 30°C, grown an additional 3 hours, and harvested at 8,000 x g. Proteins were purified at 4°C by the batch method over 1 mL Ni-NTA resin according to the manufacturer’s instructions (HisPur, Thermo Scientific). All buffers were 50 mM NaPO_4_, pH 7.4, 100 mM NaCl, with varied imidazole (10 mM lysis buffer, 20 mM wash buffer, 250 mM elution buffer). The eluted protein was dialyzed against 50 mM NaPO_4_, pH 7.4, 100 mM NaCl and either stored at 4°C or adjusted to 5% glycerol and stored at -80°C.

### Circular dichroism

CD spectroscopy to determine protein secondary structure was performed with about 5 μM protein in 50 mM NaPO_4_, pH 7.0, 50 mM NaCl as described^68^.

### L,D-transpeptidase assay with TetraRH and purified sacculi

PG sacculi for in vitro labeling were purified from *Bacillus subtilis* as outlined for *C. difficile* above, except that PG was incubated with 48% hydrofluoric acid (Sigma) instead of 6N HCl to remove teichoic acids^63^. Sacculi were adhered to poly-L-Lysine coated multiwell slides as previously described^69^. Any free poly-L-lysine coated surface was blocked with 2 mg/mL BSA for 20 min and washed with PBS. For the reaction, 5 μM enzyme was premixed with 30 μM fluorescent substrate analog in 50 mM NaPO_4_, pH 7.0, 50 mM NaCl. The reaction was started by pipetting 10 μL to the well with the PG sacculi and incubated at 37°C for 1 hour. Wells were washed with PBS and imaged by microscopy. To measure inhibition by meropenem (AuroMedics Pharma LLC, ordered from the University of Iowa hospital pharmacy), the antibiotic was added to the enzyme-TetraRh mixture, incubated 5 min at room temperature, then pipetted onto the immobilized PG sacculi and processed as above. After imaging, average fluorescence intensity was quantitated for a minimum of 10 sacculi per condition using FiJi. IC50 values were determined by performing a nonlinear fit (inhibitor vs response, 3 parameters) using GraphPad Prism v10.2.0, with the top value constrained to 100% and the bottom to 0.

### L,D-transpeptidase assay with purified disaccharide-tetrapeptide

LDTs were assayed much as described ^17^ in 50 mM NaPO_4_, pH 7.0, 50 mM NaCl at 10 µM enzyme and 30 µM disaccharide-tetrapeptide substrate (purified from R20291) in a final volume of 12 µL. As the purified substrate had been reduced prior to HPLC purification, the MurNAc was converted to the alcohol form. Assays were incubated at 37°C for 2 hours prior to chilling to 4°C and HPLC analysis using the same acetonitrile/trifluoracetic acid buffer system and peak detection at 206 nm as described above for the muropeptide analysis.

### Sequence alignments

VanW domain sequences were aligned in Clustal Omega using default parameters. Sequences were retrieved from NCBI Conserved Domains (pfam04294) using the domain boundaries defined therein. An alignment of the 35 most diverse sequences was selected, of which we chose 10 from different genera to produce an alignment that fits on one page. The sequences used are *C. difficile* R20291 Ldt4 CBE03724.1 residues 228-357, *C. difficile* R20291 Ldt5 CBE05080.1 residues 240-369, *Candidatus Desulforudis audaxviator* MP104C ACA60532.1 residues 237-362, *Moorella thermoacetica* Q2RKM6 residues 173-302, *Sulfobacillus acidophilus* DSM 10332 AEW06046.1 residues 72-201, *Ruminiclostridium cellulolyticum* H10 ACL74806.1 residues 104-233, *Gottschalkia acidurici* WP_014967910.1 residues 97-226, *Alkaliphilus oremlandii* OhILAs ABW18389.1 residues 107-236, *Syntrophomonas wolfei* Q0AX88 residues 98-227, *Desulforamulus ruminis* DSM 2154 AEG61089.1 residues 98-227, *Desulfofarcimen acetoxidans* WP_015759343 residues 119-248, *Chthonomonas calidirosea* WP_016483580.1 residues 47-176.

### Domain modeling

The VanW and YkuD domains were modeled with AlphaFold2^29^ using MMseqs2^70^ by running ColabFold v1.5.5^71^. The following parameters were used: msa_mode: mmseq2_uniref_env, pair_mode: unpaired_pair, model_type: auto, num_recycles: 3, recycle_early_stop_tolerance: auto, relax_max_iterations: 200, pairing_strategy: greedy. The highest ranked structure by pLDDT was used for illustrations. AlphFold2 structures were rendered in ChimeraX version 1.5 using default settings^72^. Structure comparison (overlay) was performed using the matchmaker function in ChimeraX using the “best-aligning pair of chains between reference and match structure”.

### Phylogenetic distribution of VanW domains

The phylogenetic distribution of VanW domains was determined by searching PF04294 in AnnoTree version 2.0 beta^39^, which includes KEGG and InterPro annotations for 80,789 bacterial and 4,416 archaeal genomes.

