## Supplementary material for "Identification of a new family of peptidoglycan transpeptidases reveals atypical crosslinking is essential for viability in *Clostridioides difficile*": Supp Figs 1-10_Supp Tables 2-4

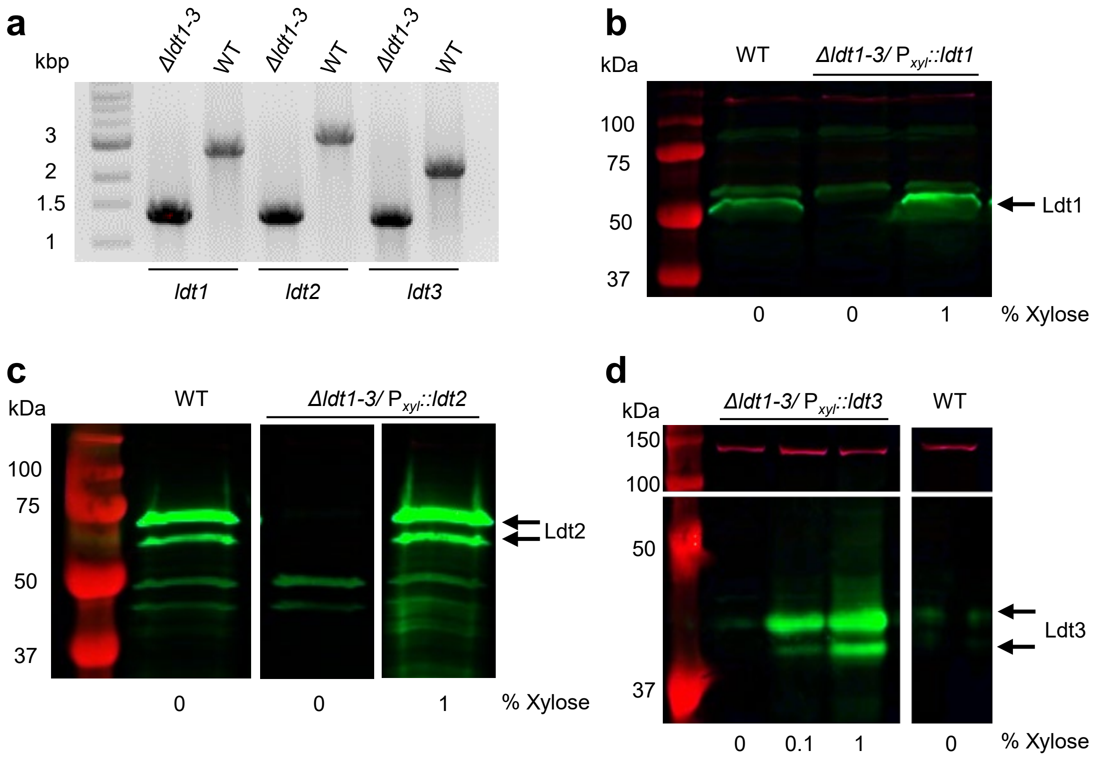

**Supplemental Fig. 1. Validation of Δ*ldt1-3* deletion mutant.** (**a**) PCR amplification products obtained using primers that flank the indicated gene deletion. Reaction mixtures were analyzed on a 1% agarose gel that was stained with ethidium bromide and photographed. Gel is representative of 2 experiments. (**b-d**) Western blots with antisera raised against soluble extracellular domain of the indicated Ldt protein. Blots are representative of at least three biological replicates. Overnight cultures of WT or the Δ*ldt1-3* mutant harboring a $P_{xyl}$::*ldt* expression plasmid were diluted 1:100 into TY or TY-thiamphenicol with or without xylose as indicated. Samples were taken for Western blotting at $OD_{600}$ ~0.8. The predicted molecular masses are: Ldt1, 52.5 kDa; Ldt2, 71.9 kDa; Ldt3, 33.3 kDa. Note that Ldt3 is essentially not expressed in WT and this blot includes an inset (above) showing an unknown biotinylated protein detected with a red streptavidin probe as a loading control. The same control is also faintly visible in the Ldt1 blot. Strains used: WT, R20291; Δ*ldt1-3*, KB124; Δ*ldt1-3*/$P_{xyl}$::*ldt1*, KB210; Δ*ldt1-3*/$P_{xyl}$::*ldt2*, KB154; and Δ*ldt1-3*/$P_{xyl}$::*ldt3*, KB181.

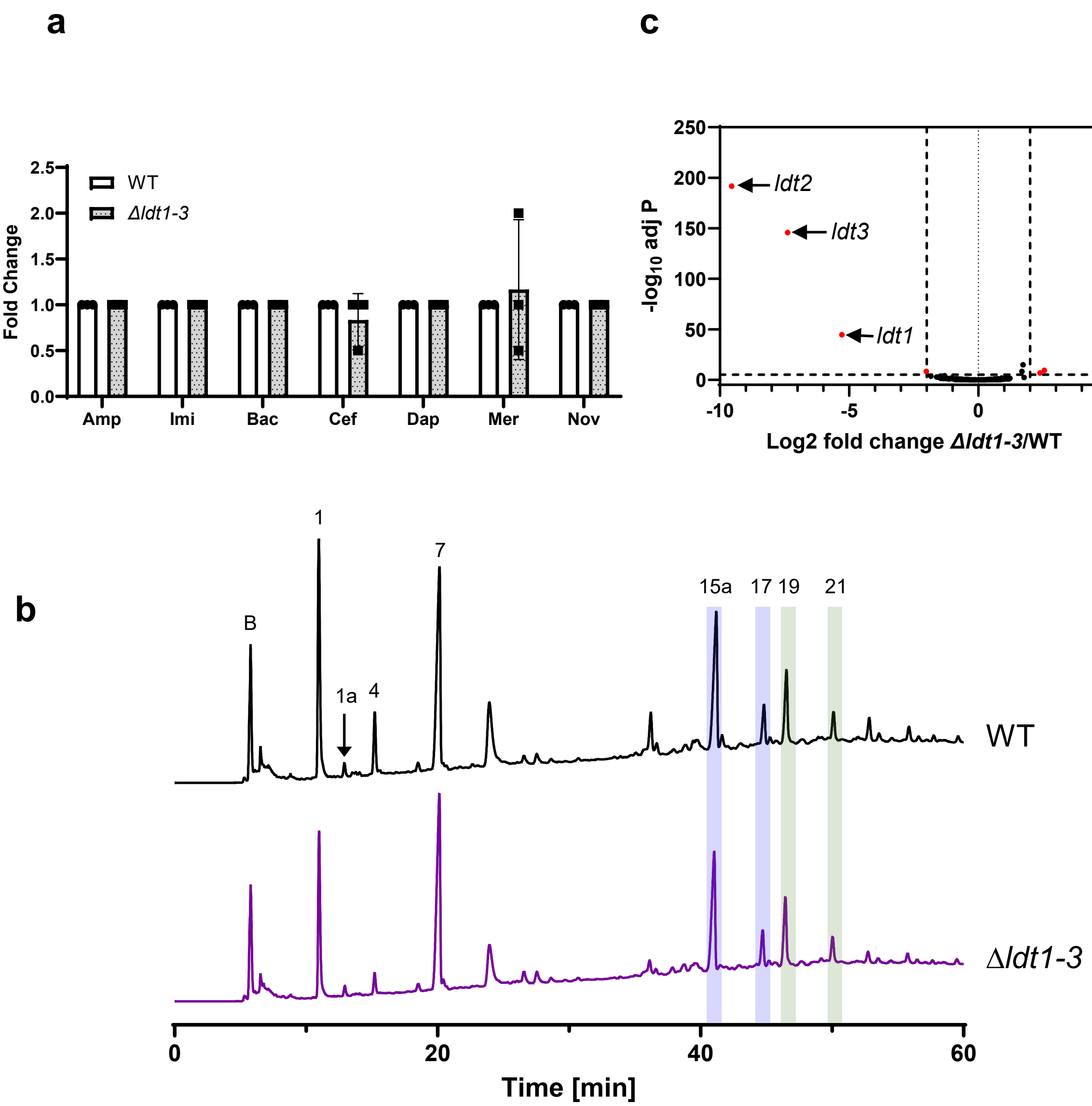

**Supplemental Fig. 2. The Δ*ldt1-3* deletion mutant is similar to wild-type in antibiotic sensitivity, transcriptional profile, and PG composition.** (**a**) Minimum inhibitory concentration of select antibiotics against the wild type (WT) strain and Δ*ldt1-3.* Data are from three biological replicates and reported as fold change difference between the strains. Abbreviations as follows with wild type MIC in brackets. Amp: ampicillin (3.1 µg/mL), Imi: imipenem (6.3 µg/mL), Bac: bacitracin (250 µg/mL), Cef: cefoxitin (200 µg/mL), Dap: daptomycin (1.6 µg/mL), Mer: meropenem (2.08 µg/mL), Nov: novobiocin (16.7 µg/mL). (**b**) Muropeptide analysis. HPLC chromatograms representative of three biological replicates are shown with the major 3-3 and 4-3 crosslinked muropeptides numbered as in Peltier et al.[15] and highlighted in blue (3-3) or green (4-3). (**c**) Volcano plot comparing transcriptome of the Δ*ldt1-3* strain to wild type growing in TY. Vertical dotted lines: log2-fold change=2; horizontal dotted line: $-\log_{10}$ adjusted P-value=5. Red dots indicate genes with log2-fold>2 and $-\log^{10}$ adjusted P-value>5. The genes encode the three deleted *ldts* and three more that just exceed the cut-offs, *cdr_2121-2122* (*sinR, sinR'*) and *cdr_2123,* a small hypothetical. Strains used: WT, R20291; Δ*ldt1-3*, KB124.

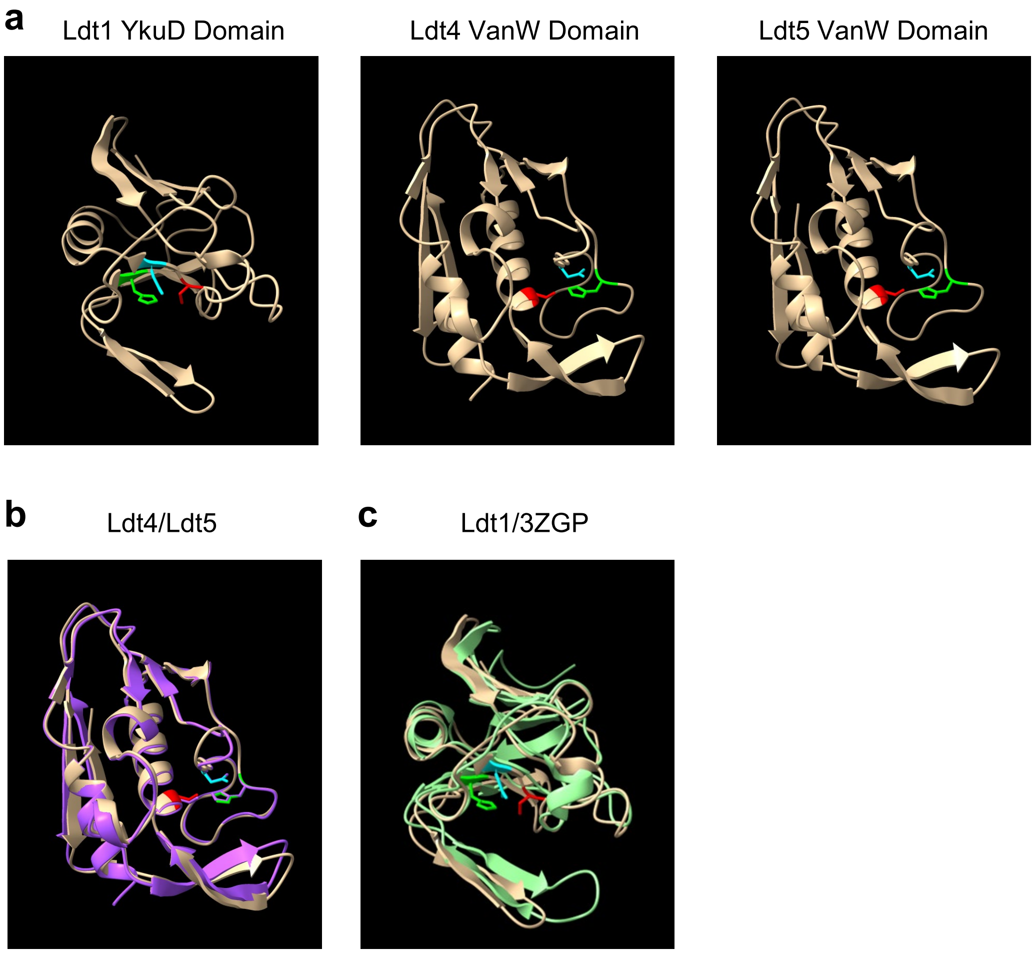

**Supplemental Fig. 3. The YkuD and VanW domains have different folds**. (**a**) AlphaFold2 structures of catalytic domains *C. difficile* Ldt1, Ldt4 and Ldt5. Catalytic triad residues are highlighted: Cys (red), His (green), Asp (cyan). (**b**) Overlay of the AlphaFold2 VanW domains from Ldt4 (purple) and Ldt5 (brown) show very similar folds. (**c**) Overlay of the AlphaFold2-predicted structure of the YkuD domain from Ldt1 (brown) with the NMR-determined structure of the YkuD domain from *E. faecium* Ldt (3ZGP, green).

```
C. difficile LDT4    NRSINIKLATDNISNVLLMPGETFSFNKHTGKRSKENGYKSAPVIMEGEMEEDYGGGVCQ
C. difficile LDT5    GRSYNVGLSARKTSDVLLMPGEEFSYNKLTGPSNKANGYKDAPVIVYGKLEQSAGGGVCQ
S. wolfei            GEETNVYIAACLLQGTVVKSGQIFSQNEKIGPYSQDKGFQPGPVYIGHQLKTTVGGGVCK
R. cellulolyticum    GEEENVHLAARLLAGTVVKPGEVFSQNNKIGPYVIARGFKKGPTYIGTKLTTTIGGGVCK
G. acidurici         GEEYNVHLAAKSLAGIVVPPGAVFSQNASIGPYTESKGYKKGPTYMGPKITTTEGGGVCK
A. oremlandii        GEEYNVHLAARTLSGTVIQPGETFSQNQRIGPYTKARGYQEGPTYIGGKVTTTEGGGVCK
D. ruminis           GEEYNIGLAASKLAGTVIKAGAVFSQNQTLGPYTQSKGYQAGPTYAGSKVTTTVGGGVCK
D. acetoxidans       AEGYNIGLAAQQLAGTVVQAEEVFSQNHTLGPYIESKGYKAGPTYSGNQLITTVGGGVCK
C. calidirosea       SQRRNARLAAWAVNGAVVPPGGLFSFDKRVGSWSADHGYVQAPVSYDGELVNAVGGGVCQ
S. acidophilus       SQAKNIELVAQRLNGTVVKPGQIFSYYARVGPYTAENGFGWGRMFVGDRIVPSIGGGVCQ
D. audaxviator       ----NAVQAAAYLNGITVQPGQVFSYNQTVGPRTAERGFVIGYAISGDRHVPARGGGVCR
M. thermoacetica     PSLHNARLAGQYLNGLVVPPGGVVSFNNVVGPRTGARGFVPGIIFMGDQKVPEIGGGICR
                         *         .  :     .*     *     .*:  .      .     ***:*:

C. difficile LDT4    VSSTLYNSVLYAGLEIVNVKNHTIPSSYVPKGRDATVADSGIDFLFKNNLKHPVYIKNYV
C. difficile LDT5    TSSTVYNAALLSGMEITQVTNHSSASTYVPKGRDATVSDGGLNLKFKNPYKHPVYIKNYA
S. wolfei            IASTLYNVTILSNLPVIERYAHSMPVPYVPLGQDATVCYGVKDFKFLNNSPYPILIWAES
R. cellulolyticum    MASTLYNVAILSNLPVVERHAHSMPVPYVPYGQDATVSYGNKDLKFKNDTSSPIMIWAQG
G. acidurici         IASTLYNVAIYSNLEVVERYNHTMPVPYVPYGQDATVAYGFKDLKFKNNTDFPILIWAEG
A. oremlandii        IASTLYNVAILSDLQIVERHNHGMPVPYVPYGQDATVAYGAKDIRFKNNTDSPILIWSVG
D. ruminis           IASLLYNVATLSDLQIIMRYPHSMTVPYVPPGQDATVFCGVKDLRFLNDTGGSVMIWSQK
D. acetoxidans       IASMLYNVVTFCDLKVISRSPHSMTVPYVPPGQDATVYYGCRDFSFFNDSGRPILIWAQK
C. calidirosea       TSSTLYNAALLAGMQIVERHPHHFCPEYVPPGRDAAVAQTIIDLRFKNPYPWPVRIECRA
S. acidophilus       GSSTLYAALLRTGLPIIERHHHGLTVPYLPPGEDATVASDYLDFRFKNNRTTPILITAQA
D. audaxviator       TSTVLYGAVLNAGLPVIERHAHTRPVGYVPMGRDATVSYGTADLKFRNDLPRPVRIKAGG
M. thermoacetica     TATLLHNAVLSAGLEVVERHRHGLPVTYVPPGYDATVYYGVLDYRFRNNRPVPIKLEFTS
                      :: ::      .: :     *     *:* * **:*     :  * *     : :

C. difficile LDT4    SGNQIVCNIY
C. difficile LDT5    GGGSVSSVIY
S. wolfei            IGNRLYIAFY
R. cellulolyticum    VDNILYVAFY
G. acidurici         IENRLYIGFY
A. oremlandii        IDNTLYIGFY
D. ruminis           VGNTVYMALY
D. acetoxidans       EGDTLYMAFY
C. calidirosea       TQDTLEASFW
S. acidophilus       GQRHLTVAIW
D. audaxviator       TVRQLQVTLW
M. thermoacetica     QGSSITMAIW
                         :   ::
```

**Supplemental Fig. 4. VanW domain amino acid sequence alignment**. Proposed catalytic triad is highlighted with red, green and blue. Gray highlight and asterisks denote strict amino acid identity, colons and periods indicate other conserved positions. Sequences shown are from *C. diffficile*, *Desulforudis audaxviator, Moorella thermoacetica, Sulfobacillus acidophilus, Ruminiclostridium cellulolyticum, Gottschalkia acidurici, Alkaliphilus oremlandii, Syntrophomonas wolfei, Desulforamulus ruminis, Desulfofarcimen acetoxidans*, and *Chthonomonas calidirosea*.

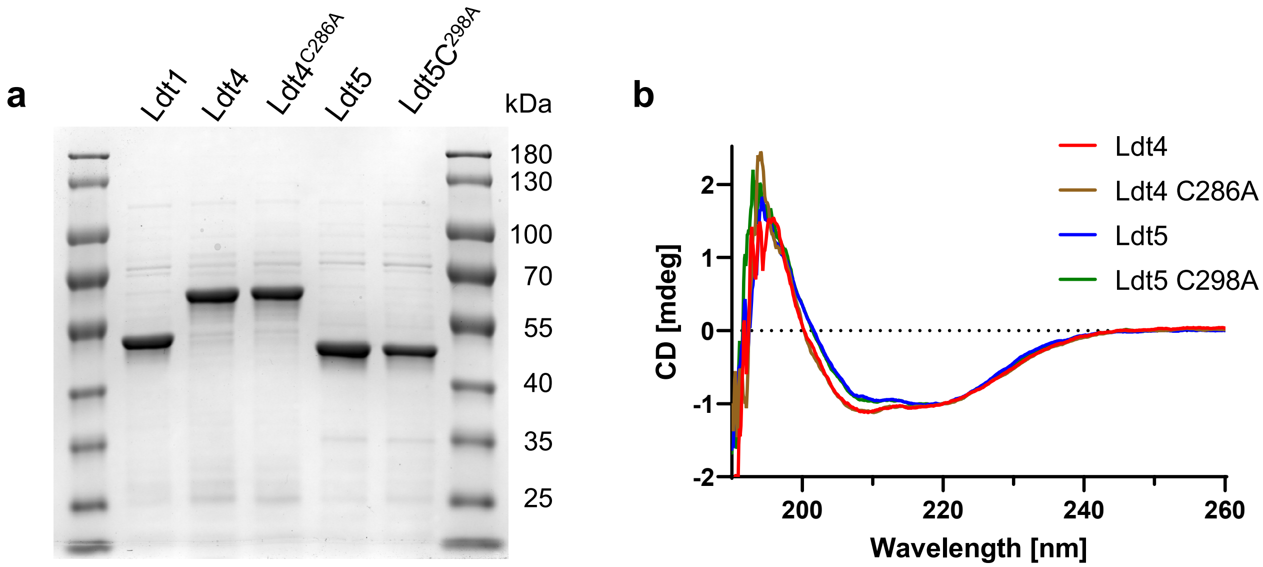

**Supplemental Fig. 5. Purified Ldts.** His-tagged Ldts without the N-terminal transmembrane region were expressed in *E. coli* and purified over a Nickel-affinity resin. (**a**) Purity determination by SDS-PAGE. About 4 µg of purified protein were separated by 10% SDS-PAGE and stained with Coomassie Blue. Molecular mass standards are indicated to the right of the gel. Predicted molecular weights: Ldt1, 51 kD; Ldt4 54 kD; Ldt5, 45 kD. (**b**) Circular dichromism (CD) spectroscopy to evaluate protein folding. CD spectra were generated for 5 µM enzyme and normalized to the absorbance at 220 nm. Ldts with the active site mutation showed the same degree of folding as the corresponding wild type enzyme.

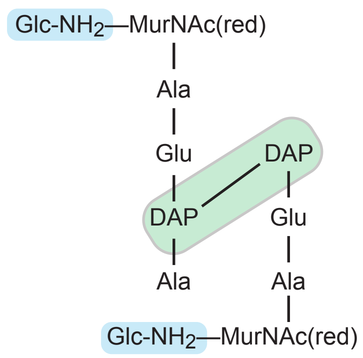

| Fragment | $[M+H]^+$ | $[M+H - H_2O]^+$ | $[M+H - 2H_2O]^+$ |
|---|---|---|---|
| $(DAP)_2$ | 363.1874 | 345.1769 | |
| $(Glu)(DAP)_2$ | 492.2300 | 474.2194 | |
| $(Glu)_2(DAP)_2$ | 621.2726 | 603.2620 | |
| $Ala(Glu)_2(DAP)_2$ | 692.3097 | 674.2992 | |
| $MurNAc(Ala)(Glu)(DAP)_2$ | 840.3833 | 822.3727 | |
| $MurNAc(Ala)_2(Glu)(DAP)_2$ | 911.4204 | 893.4098 | |
| $MurNAc(Ala)(Glu)_2(DAP)_2$ | 969.4259 | 951.4153 | 933.4076 |

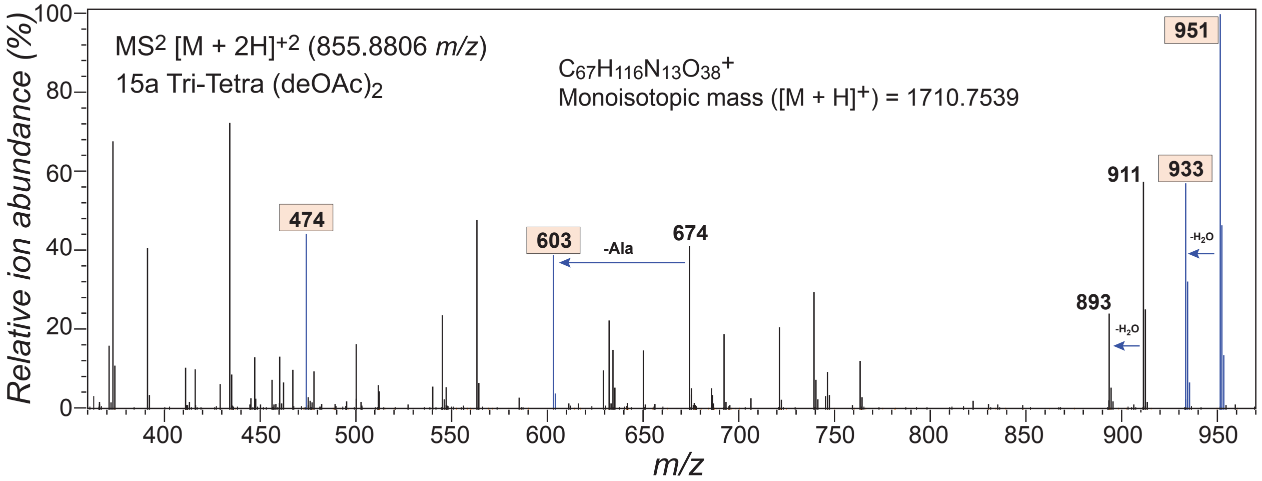

**Supplemental Fig. 6.** *Structural analysis of the predominant Ldt4 and Ldt5 reaction product.* Confirmation of the 3-3 crosslink was determined by manual interpretation of fragmentation data that solely supported DAP-DAP bonds (beige ions). The ions indicated in gray were present but not indicative of a 3-3 crosslink. While the DAP dimer (363 *m/z*) was observed, the ion intensity was too weak to be used for confirmation. However, ions associate with Glu and DAP (474, 603 *m/z*) fully supported the stucture shown along several ions containing an intact NAM(red) moiety and only one alanine (951, 933 *m/z*).

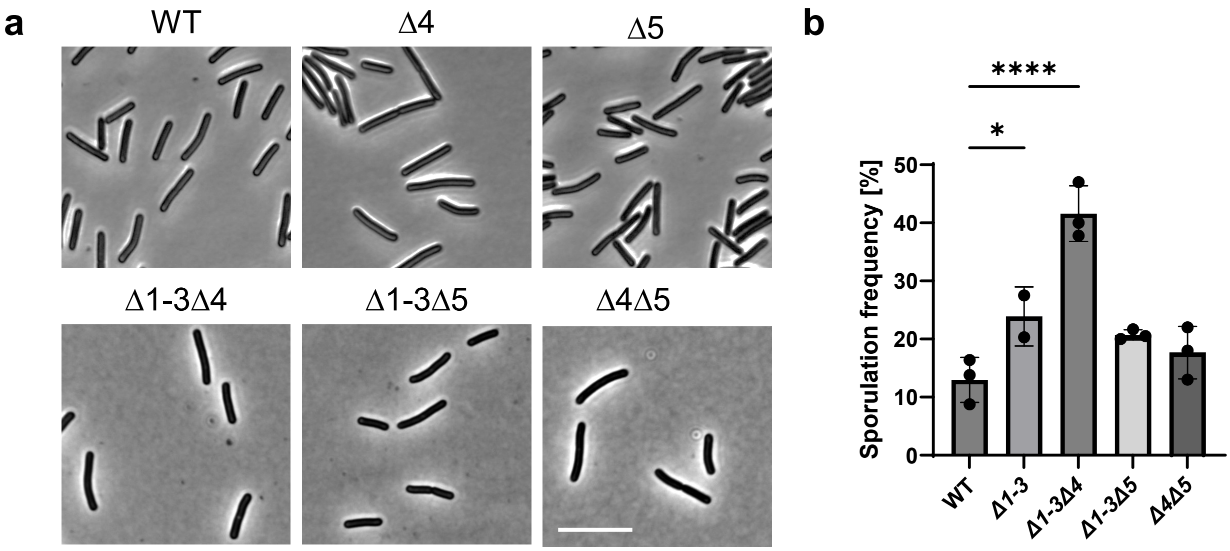

**Supplemental Fig. 7: Morphology and sporulation of *ldt* deletion mutants.** (**a**) Phase contrast images of strains grown to mid-log. Size bar: 10 µm. Images are representative of at least three biological replicates. (**b**) Sporulation efficiency. Dots indicate the values from two or three biological replicates. Bars and error bars indicate the mean ± s.d. *, $p < 0.05$. **** $p < 0.001$ in a two-way ANOVA. Frequency is calculated as the number of spores divided by the sum of spores plus vegetative cells [spores/(spores + vegetative cells)]. Strains shown: WT, R20291; Δ4, KB439; Δ5, KB440; Δ1-3Δ4, KB474 ; Δ1-3Δ5, KB502; and Δ4Δ5, KB529.

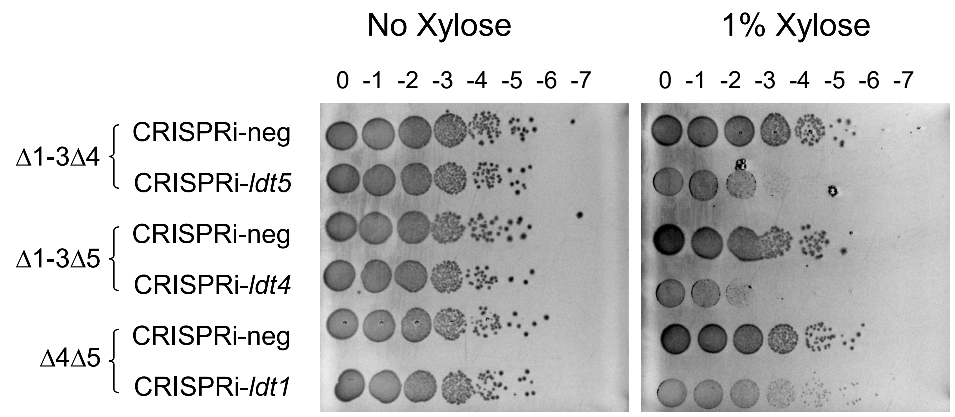

**Supplemental Fig. 8: Ldt1, Ldt4, or Ldt5 is sufficient for viability and Ldt2 plus Ldt3 are not sufficient.** CRISPRi plasmids targeting *ldt5, ldt4, or ldt1* were introduced into the indicated deletion strains. The negative control plasmid contained sgRNA against sequence not found in *C. difficile.* Serial dilutions of overnight cultures were spotted onto TY plates with or without 1% xylose, and plates were imaged after overnight incubation. Images are representative of three biological replicates. Strains shown: Δ1-3Δ4/neg, KB579; Δ1-3Δ4/CRISPRi-*ldt5*, KB508; Δ1-3Δ5/neg, KB580; Δ1-3Δ5/CRISPRi-*ldt4*, KB514 ; Δ4Δ5/neg, KB565; and Δ4Δ5/CRISPRi-*ldt1*, KB566

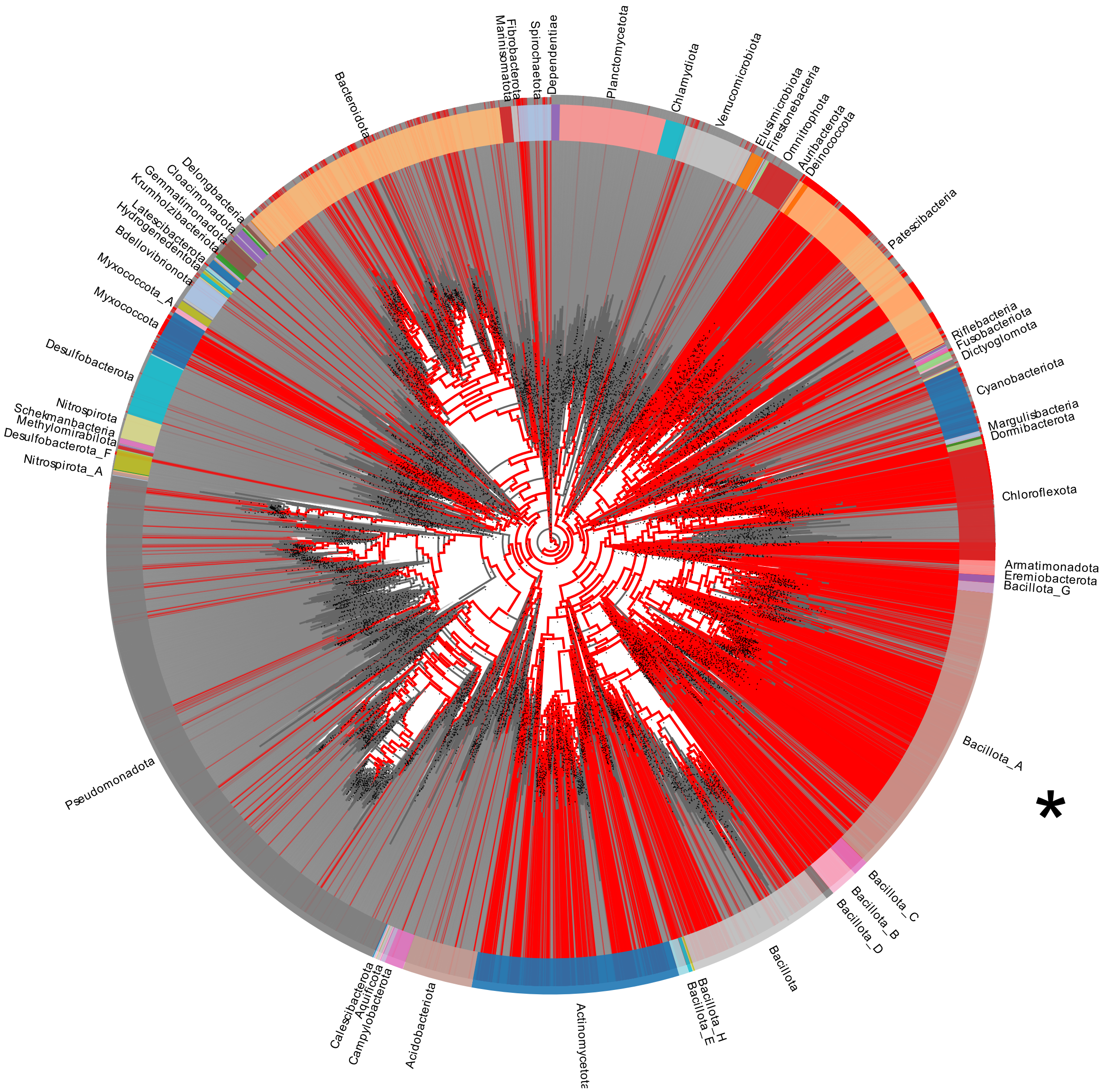

**Supplemental Fig. 9:** Taxonomic distribution of VanW domains. Red lines indicate the presence of VanW domais, gray lines the absence. Outer ring indicates bacterial phyla at the level of Class. *C. difficile* belongs to Bacillota A (asterisk). Some phyla names were pruned for clarity. Figure prepared using AnnoTree v1.3 Beta.

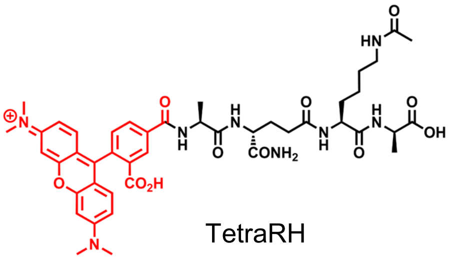

TetraRH

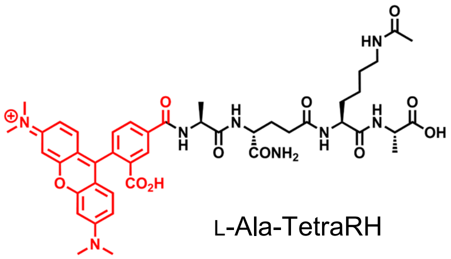

L-Ala-TetraRH

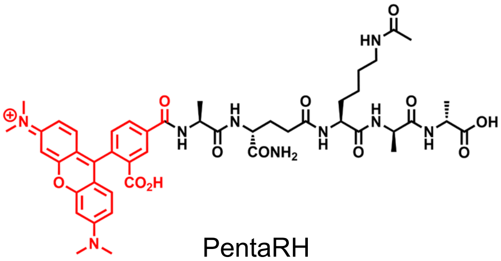

PentaRH

**Supplemental Fig. 10. Fluorescent substrate analogs**. TetraRH: L,D-transpeptidase specific substrate analog, Rhodamine-L-Ala-iso-D-Gln-L-Lys(Ac)-D-Ala. L-Ala-TetraRH: negative control, Rhodamine-L-Ala-iso-D-Gln-L-Lys(Ac)-L-Ala; PentaRH: PBP specific substrate analog, Rhodamine-L-Ala-iso-D-Gln-L-Lys(Ac)-D-Ala-D-Ala.

Supplemental Table 2. Strains used in this study.

| **Strain and species** | **Genotype and/or description** | **Alternate name** | **Source or reference** |
|---|---|---|---|
| *E. coli* | | | |
| OmniMAX-2T1R | F′ [*proAB+ lacI* q *lacZΔ* M15 Tn*10* (Tetr) *Δ(ccdAB* )] *mcrA Δ* (*mrr-hsdRMS-mcrBC* ) φ80(*lacZ* )*Δ* M15 *Δ* (*lacZYA-argF* )*U169 endA1 recA1 supE44 thi-1 gyrA96 relA1 tonA panD* | | Invitrogen |
| HB101/pRK24 | F− *mcrB mrr hsdS20* (rB− mB−) *recA13 leuB6 ara* − *14 proA2 lacY1 galK2 xyl-5 mtl-1 rpsL20* | | (1) |
| MG1655 | Wild type isolate | | (2) |
| Rosetta(DE3) | $F^-$ *ompT hsdS* $_B$($r_B^-$ $m_B^-$) *gal dcm* (DE3) pRARE ($Cam^R$) | | Sigma-Aldrich |
| KB005 | Rosetta(DE3)/pKB001 | | This study |
| KB007 | Rosetta(DE3)/pKB003 | | This study |
| KB008 | Rosetta(DE3)/pKB004 | | This study |
| CE4771 | Rosetta(DE3)/pCE1169 | | This study |
| CE4772 | Rosetta(DE3)/pCE1170 | | This study |
| CE4777 | Rosetta(DE3)/pCE1173 | | This study |
| CE4778 | Rosetta(DE3)/pCE1174 | | This study |
| *B. subtilis* | | | |
| BS49 | Tn*916* donor strain, $Tet^r$ | | (3) |
| PY79 | Wild type strain | | (4) |
| *C. difficile* , all R20291 or derivatives | | | |
| R20291 | Wild type strain from UK outbreak (ribotype 027) | | (5) |
| KB071 | *Δldt1* | | This study |
| KB075 | *Δldt2* | | This study |
| KB130 | *Δldt3* | | This study |
| KB103 | *Δldt1Δldt2* | | This study |
| KB139 | *Δldt1Δldt3* | | This study |
| KB166 | *Δldt2Δldt3* | | This study |
| KB124 | *Δldt1Δldt2Δldt3* | Δ1-3 | This study |
| KB464 | R20291/pBZ101 | | This study |
| KB465 | Δ1-3/pBZ101 | | This study |
| KB154 | Δ1-3/pCE0938 | | This study |
| KB181 | Δ1-3/pCE0983 | | This study |
| KB210 | Δ1-3/pKB025 | | This study |
| KB439 | Δ*ldt4* | Δ4 | This study |
| KB440 | Δ*ldt5* | Δ5 | This study |
| KB474 | Δldt1Δldt2Δldt3Δ*ldt4* | Δ1-3Δ4 | This study |
| KB502 | Δldt1Δldt2Δldt3Δ*ldt5* | Δ1-3Δ5 | This study |
| KB529 | Δ*ldt4* Δ*ldt5* | Δ4Δ5 | This study |
| KB547 | Δ*ldt1-3*Δ*ldt4* $P_{tet}$::*ldt5* | Δ1-3Δ4 $P_{tet}$::ldt5 | This study |
| KB548 | Δ1-3Δ4 $P_{tet}$::*ldt5*/pBZ101 | | This study |
| KB549 | Δ1-3Δ4 $P_{tet}$::*ldt5*/pKB025 | | This study |
| KB550 | Δ1-3Δ4 $P_{tet}$::*ldt5*/pCE0938 | | This study |
| KB551 | Δ1-3Δ4 $P_{tet}$::*ldt5*/pCE0983 | | This study |
| KB552 | Δ1-3Δ4 $P_{tet}$::*ldt5*/pCE1175 | | This study |
| KB553 | Δ1-3Δ4 $P_{tet}$::*ldt5*/pCE1176 | | This study |

Supplemental Table 2. Strains used in this study.

| Strain and species | Genotype and/or description | Alternate name | Source or reference |
|---|---|---|---|
| KB508 | Δ1-3Δ4/pKB081 | | This study |
| KB514 | Δ1-3Δ5/pKB083 | | This study |
| KB566 | Δ4Δ5/pIA68 | | This study |
| KB565 | Δ4Δ5/pIA34 | | This study |
| KB579 | Δ1-3Δ4/pIA34 | | This study |
| KB580 | Δ1-3Δ5/pIA34 | | This study |
| KB633 | Δ1-3Δ4/pBZ101 | | This study |
| KB634 | Δ1-3Δ4/pCE1176 | | This study |
| KB635 | Δ1-3Δ4/pCE1186 | | This study |
| KB636 | Δ1-3Δ4/pCE1191 | | This study |
| KB637 | Δ1-3Δ4/pCE1198 | | This study |
| KB638 | Δ1-3Δ4/pCE1199 | | This study |
| KB639 | Δ1-3Δ4/pCE1200 | | This study |
| KB640 | Δ1-3Δ4/pCE1201 | | This study |
| KB641 | Δ1-3Δ4/pCE1202 | | This study |

Supplemental Table 3. Plasmids used in this study.

| Plasmid | Relevant features | Parent vector | Restriction enzymes to digest parent vector | PCR primers | PCR template | Assembly | Comments | Reference |
|---|---|---|---|---|---|---|---|---|
| pAP114 | $P_{xyl}$::*mCherryOpt catP* | | | | | | Parent plasmid for cloning under $P_{xyl}$ control | (6) |
| pBZ101 | $P_{xyl}$ empty vector *catP* | | | | | | Empty vector control for $P_{xyl}$ induction | (7) |
| pCE678 | $P_{xyl}$::*Cas9-opt* $P_{gdh}$::*sgRNA-pgdA-2,* homology to delete *pgdA catP* | | | | | | Parent plasmid for CRISPR editing | (8) |
| pCE938 | $P_{xyl}$::*ldt2 catP* | pAP114 | BamHI, SacI | 5609+5610 | R20291 | ITA[1] | *cdr_2601 (ldt2)* from *C. difficile* R20291 under xylose control | This study |
| pCE983 | $P_{xyl}$::*ldt3 catP* | pAP114 | BamHI, SacI | 6243+5612 | R20291 | ITA | *cdr_2843 (ldt3)* from *C. difficile* R20291 under xylose control | This study |
| pCE1169 | $P_{tac}$::*6xHis-ldt4*$^{28-489}$ *ampR* | pET21a-6xHis-rTEV | NcoI, EcoRI | 6875+6876 | R20291 | ITA | Expression plasmid: N-terminal His tagged Ldt4$^{28-489}$, no transmembrane region | This study |
| pCE1170 | $P_{tac}$:: *6xHis-ldt5*$^{38-419}$ *ampR* | pET21a-6xHis-rTEV | NcoI, EcoRI | 6878+6879 | R20291 | ITA | Expression plasmid: N-terminal His tagged Ldt5$^{38-419}$, no transmembrane region | This study |
| pCE1172 | $P_{xyl}$::*Cas9* $P_{gdh}$::*sgRNA-cdr_985, homology region to replace* $P_{ldt5}$ with $P_{tet}$, *catP* | pIA123 | PstI | 6886+6887<br>6888+6889<br>6890+6891 | R20291<br>R20291<br>pRPF185 | ITA | Plasmid intermediate: inserts homology regions to replace $P_{ldt5}$ with $P_{tet}$ into CRISPR editing plasmid pIA123 | This study |
| pCE1173 | $P_{tac}$::*6xHis-ldt4*$^{28-489}$ *C286A ampR* | pET21a-6xHis-rTEV | NcoI, EcoRI | 6875+6897<br>6896+6876 | R20291<br>R20291 | ITA | Expression plasmid: N-terminal His tagged Ldt4$^{28-489}$ catalytic mutant C286A, no transmembrane region | This study |
| pCE1174 | $P_{tac}$::*6xHis-ldt5*$^{38-419}$ *C298A ampR* | pET21a-6xHis-rTEV | NcoI, EcoRI | 6878+6899<br>6898+6879 | R20291<br>R20291 | ITA | Expression plasmid: N-terminal His tagged Ldt5$^{38-419}$ catalytic mutant C298A, no transmembrane region | This study |
| pCE1175 | $P_{xyl}$::*ldt4 catP* | pAP114 | BamHI, SacI | 6892+6893 | R20291 | ITA | *cdr_1285 (ldt4)* from *C. difficile* R20291 under xylose control | This study |
| pCE1176 | $P_{xyl}$::*ldt5 catP* | pAP114 | BamHI, SacI | 6894+6895 | R20291 | ITA | *cdr_2055 (ldt5)* from *C. difficile* R20291 under xylose control | This study |
| pCE1180 | $P_{xyl}$::*Cas9* $P_{gdh}$::*sgRNA-ldt5,* homology region to replace $P_{ldt5}$ with $P_{tet}$, *catP* | pCE1172 | MscI, MluI | 6906+4237 | pIA33 | ITA | Vector suitable for CRISPR replacing $P_{ldt5}$ with $P_{tet}$; replaces *sgRNA-cdr_985* with *sgRNA-ldt5* in pCE1172 | This study |
| pCE1186 | $P_{xyl}$::*yoaR* | pAP114 | BamHI, SacI | 6922+6923 | *Bacillus subtilis* PY79 | ITA | *yoaR* from *Bacillus subtilis* PY79 under xylose control | This study |
| pCE1191 | $P_{xyl}$::*vanW*$_{Ef}$ | pAP114 | BamHI, SacI | 6932+6933 | *Enterococcus faecalis* V583 | ITA | *vanW* from *Enterococcus faecalis* V583 under xylose control | This study |
| pCE1198 | $P_{xyl}$::*vanW*$_{Pb}$ | pAP114 | BamHI, SacI | 7013+7014 | *Paraclostridium bifermentans* 638 | ITA | *vanW* (WP_021433578) from *Paraclostridium bifermentans* 638 under xylose control | This study |
| pCE1199 | $P_{xyl}$::*vanW*$_{Rum}$ | pAP114 | BamHI, SacI | 7015+7016 | *Ruminococcaceae bacterium* D16 | ITA | *vanW* (HMPREF0866_01899) from *Ruminococcaeae bacterium* D16 under xylose control | This study |
| pCE1200 | $P_{xyl}$::*vanW*$_{Lac}$ | pAP114 | BamHI, SacI | 7017+7018 | *Lachnospiraceae bacterium* 5_1_57FAA | ITA | *vanW* (HMPREF0993_00855) from *Lachnospiraceae bacterium* 5_1_57FAA under xylose control | This study |
| pCE1201 | $P_{xyl}$::*vanW*$_{Pep0521}$ | pAP114 | BamHI, SacI | 7021+7022 | *Peptostreptococcaceae bacterium* AS15 | ITA | *vanW* (HMPREF1142_0521) from *Peptostreptococcaceae bacterium* AS15 | This study |
| pCE1202 | $P_{xyl}$::*vanW*$_{Pep1713}$ | pAP114 | BamHI, SacI | 7023+7024 | *Peptostreptococcaceae bacterium* AS15 | ITA | *vanW* (HMPREF1142_1713) from *Peptostreptococcaceae bacterium* AS15 | This study |
| pET21a-6xHIS-rTEV | | | | | | | pET21a vector (Novagen) with N-terminal His6 tag and tobacco etch virus (rTEV) protease cleavage site | (9) |
| pCI5492 | $P_{xyl}$::*dCas9-opt* $P_{gdh}$::*sgRNA-cdr_985* | pIA33 | MscI, NotI | 5492+4084 | pIA33 | ITA | CRISPRi against *cdr_985 (pbp2)* | This Study |
| pIA33 | $P_{xyl}$::*dCas9-opt* $P_{gdh}$::*sgRNA-rfp catP* | | | | | | Parent plasmid for CRISPRi constructs | (6) |
| pIA34 | $P_{xyl}$::*dCas9-opt* $P_{gdh}$::*gRNA-neg* | | | | | | CRISPRi negative control plasmid | (6) |

Supplemental Table 3. Plasmids used in this study.

| Plasmid | Relevant features | Parent vector | Restriction enzymes to digest parent vector | PCR primers | PCR template | Assembly | Comments | Reference |
|---|---|---|---|---|---|---|---|---|
| pIA68 | $P_{xyl}$::*dCas9-opt* $P_{gdh}$::*sgRNA-ldt1* | pIA34 | MscI, MluI | | pIA33 | ITA | CRISPRi against *ldt1* | This study |
| pIA123 | $P_{xyl}$::*Cas9* $P_{gdh}$::*sgRNA-cdr_985 catP* | pCI5492 | SalI, XhoI | 6269+6270 | pCE678 | ITA | CRISPR editing parent plasmid; sgRNA-*cdr_985 (pbp2)*; no homology region | This study |
| pKB001 | $P_{tac}$::6xHis-*ldt1*$^{38-469}$ *ampR* | pET21a-6xHis-rTEV | NcoI, EcoRI | 5467+5468 | R20291 | ITA | Expression plasmid: N-terminal 6x His tag on Ldt1$^{38-469}$, no transmembrane section | This study |
| pKB003 | $P_{tac}$::6xHis-*ldt2*$^{38-617}$ *ampR* | pET21a-6xHis-rTEV | NcoI, EcoRI | 5471+5472 | R20291 | ITA | Expression plasmid: N-terminal His tagged Ldt2$^{38-617}$, no transmembrane region | This study |
| pKB004 | $P_{tac}$::*6xHis-ldt3*$^{7-289}$ *ampR* | pET21a-6xHis-rTEV | NcoI, EcoRI | 5473+5474 | R20291 | ITA | Expression plasmid: N-terminal His tagged Ldt3$^{7-289}$, no transmembrane region | This study |
| pKB007 | $P_{xyl}$::*Cas9-opt* $P_{gdh}$::*sgRNA-pgdA-2,* homology to delete *ldt2 catP* | pCE678 | NotI, XhoI | 5541+5542, 5543+5544 | R20291 R20291 | ITA | Plasmid intermediate to build *ldt2* deletion plasmid | This study |
| pKB009 | $P_{xyl}$::*Cas9-opt* $P_{gdh}$::*sgRNA-pgdA-2,* homology to delete *ldt3 catP* | pCE678 | NotI, XhoI | 5463+5464, 5465+5466 | R20291 R20291 | ITA | Plasmid intermediate to build *ldt3* deletion plasmid | This study |
| pKB015 | $P_{xyl}$::*Cas9-opt* $P_{gdh}$::*sgRNA-pgdA-2,* homology to delete *ldt1 catP* | pCE678 | NotI, XhoI | 5459+5460, 5461+5462 | R20291 R20291 | ITA | Plasmid intermediate to build *ldt1* deletion plasmid | This study |
| pKB019 | $P_{xyl}$::*Cas9-opt* $P_{gdh}$::*sgRNA-ldt1,* homology to delete *ldt1 catP* | pKB15 | MscI, MluI | 5448+4237 | pCE678 | ITA | CRISPR edit plasmid to delete *ldt1* | This study |
| pKB022 | $P_{xyl}$::*Cas9-opt* $P_{gdh}$::*sgRNA-ldt2,* homology to delete *ldt2 catP* | pKB07 | MscI, MluI | 5540+4237 | pCE678 | ITA | CRISPR edit plasmid to delete *ldt2* | This study |
| pKB024 | $P_{xyl}$::*Cas9-opt* $P_{gdh}$::*sgRNA-ldt3,* homology to delete *ldt3 catP* | pKB09 | MscI, MluI | 5456+4237 | pCE678 | ITA | CRISPR edit plasmid to delete *ldt3* | This study |
| pKB025 | $P_{xyl}$::*ldt1 catP* | pAP114 | BamHI, SacI | 5607+5608 | R20291 | ITA | *cdr_2797 (ldt1)* from *C. difficile* R20291 under xylose control | This study |
| pKB067 | $P_{xyl}$::*Cas9* $P_{gdh}$::*sgRNA-cdr_985,* homology to delete *ldt5 catP* | pIA123 | PstI | 6737+6738, 6739+6740 | R20291 R20291 | ITA | Plasmid intermediate to build *ldt5* deletion plasmid | This study |
| pKB068 | $P_{xyl}$::*Cas9* $P_{gdh}$::*sgRNA-cdr_985,* homology to delete *ldt4 catP* | pIA123 | PstI | 6733+6734, 6735+6736 | R20291 R20291 | ITA | Plasmid intermediate to build *ldt4* deletion plasmid | This study |
| pKB071 | $P_{xyl}$::*Cas9* $P_{gdh}$::*sgRNA-ldt5,* homology to delete *ldt5 catP* | pKB67 | MscI, MluI | 6780+4237 | pIA123 | ITA | CRISPR edit plasmid to delete *ldt5* | This study |
| pKB073 | $P_{xyl}$::*Cas9* $P_{gdh}$::*sgRNA-ldt4,* homology to delete *ldt4 catP* | pKB68 | MscI, MluI | 6774+4237 | pIA123 | ITA | CRISPR edit plasmid to delete *ldt4* | This study |
| pKB081 | $P_{xyl}$::*dCas9-opt* $P_{gdh}$::*sgRNA-ldt5* | pIA34 | MscI, MluI | 6865+4237 | pIA34 | ITA | CRISPRi against *ldt5* | This study |
| pKB083 | $P_{xyl}$::*dCas9-opt* $P_{gdh}$::*sgRNA-ldt4* | pIA34 | MscI, MluI | 6871+4237 | pIA34 | ITA | CRISPRi against *ldt4* | This study |
| pRPF185 | $P_{tet}$::*gusA* | | | | | | Source of $P_{tet}$ | (10) |

[1]: Isothermal assembly

Supplemental Table 4. Oligonucleotides used in this study.

| Oligo | Sequence | Use |
|---|---|---|
| 4084 | AACTTATAGGATCCGCGGCCGCTAGTCAGACATCATGCTGATCTAGA | Cloning |
| 4237 | CTTATAGGATCCGCGGCCGCTAG | Cloning |
| 5448 | AATTAAACTGTAAATGGCCAAATTGTAATATCTTTACCTG GTTTTAGAGCTAGAAATAGC | Cloning |
| 5456 | AATTAAACTGTAAATGGCCAATTTTTAAGAAAGATAATGG GTTTTAGAGCTAGAAATAGC | Cloning |
| 5459 | AAACAGCTATGACCGCGGCCGCGTTGAAGACATTACGAAACTAG | Cloning |
| 5460 | AACTGTTAGCAACACATTTAAATTAAATCCTTCCTTACATTG | Cloning |
| 5461 | ATGTAAGGAAGGATTTAATTTAAATGTGTTGCTAACAGTT | Cloning |
| 5462 | TTATTTTTATGCTAGCTCGAGCCTCATTGTTAAAGTATAAACA | Cloning |
| 5463 | AAACAGCTATGACCGCGGCCGCTTAAAAGGTGAAATAATCTGT | Cloning |
| 5466 | TTATTTTTATGCTAGCTCGAGAGACTATGAAGGTATCAAC | Cloning |
| 5467 | TGTATTTTCAGGGCGCCATGAGAAATCATTTTTACTTTGGA | Cloning |
| 5468 | TCGACGTAGGCCTTTGAATTCTAGTATAAAATAATTGGTGTACC | Cloning |
| 5471 | TGTATTTTCAGGGCGCCATGAGTAAACATGTGATTATAGTAAA | Cloning |
| 5472 | TCGACGTAGGCCTTTGAATTCTATTTTGCAAGAATATCCA | Cloning |
| 5473 | TGTATTTTCAGGGCGCCATGAAATTAACTAAACTAAATATTAAAAAATATA | Cloning |
| 5474 | TCGACGTAGGCCTTTGAATTTTAATGAATTATAACTGTTGTTG | Cloning |
| 5492 | AATTAAACTGTAAATGGCCACTGCTATTGAAACACCAACA GTTTTAGAGCTAGAAATAGC | Cloning |
| 5540 | AATTAAACTGTAAATGGCCACCTATAATTGTATCCCATGT GTTTTAGAGCTAGAAATAGC | Cloning |
| 5541 | AAACAGCTATGACCGCGGCCGCAACTCAATAGTGGTTGAT | Cloning |
| 5542 | TTTTTCAATATATATTTTTATACTACATGTGTCTAATTATAACAT | Cloning |
| 5543 | ATAATTAGACACATGTAGTATAAAAATATATATTGAAAATAATTATAATTGAG | Cloning |
| 5544 | TTATTTTTATGCTAGCTCGAGGCTCTACAATAGGAACTTC | Cloning |
| 5607 | CGATAGTTATGAAGTGAGCTGTAAGGAAGGATTTAATTATGATTGATG | Cloning |
| 5608 | TTATTAAAACTTATAGGATCCTTCGCTTAACTGTTAGCAAC | Cloning |
| 5609 | CGATAGTTATGAAGTGAGCTTAAGGAGGATGTAGTAATGTTTCTAAAAGAGGGGGA | Cloning |
| 5610 | TTTTATTAAAACTTATAGGATCCACATCTCAATTATAATTATTTTTCAATATATATTTTTACTATTTTGCA | Cloning |
| 5612 | AGTTTTATTAAAACTTATAGGATCTTGAGAATCATTATCCCAATAAAAACTC | Cloning |
| 6243 | AGATACCATAGATCCGGTACCATGAATTATAACTGTTGTTGTATCTG | Cloning |
| 6269 | AGGAGGGTAAAGAGGAGAGTCGACGCATGGATAAAAAATATAGTATAGGATTAGATATAG | Cloning |
| 6270 | CCGTATTTCTACGATGTTTTTGCTCGAGTTAATCACCACCTAATTGAGATAAA | Cloning |
| 6733 | CAGGAAGGGCGAATTCTGCATGGTAAGTGCAAAAACTAAA | Cloning |
| 6736 | GAGACCGGTCAGATCTGCACCTTGTTTATAAAGTTCATCTAGT | Cloning |
| 6737 | CAGGAAGGGCGAATTCTGCAAAAACAATTAGACGAAATAATAGA | Cloning |
| 6738 | ATATTTCTCTAAAAAATTATAACTTCACCTCATTTTGACA | Cloning |
| 6739 | TGTCAAAATGAGGTGAAGTTATAATTTTTTAGAGAAATATTTGTAATAT | Cloning |
| 6740 | GAGACCGGTCAGATCTGCACACATCTATTATTTTTAGTATATAAGGA | Cloning |
| 6774 | AATTAAACTGTAAATGGCCAAATGTGCATGCAATTTAAAG GTTTTAGAGCTAGAAATAGC | Cloning |
| 6780 | AATTAAACTGTAAATGGCCATTGTATGAAAATTCTTCACC GTTTTAGAGCTAGAAATAGC | Cloning |
| 6865 | AATTAAACTGTAAATGGCCATAAGTAAGATTTAAGTCTTC GTTTTAGAGCTAGAAATAGC | Cloning |
| 6871 | AATTAAACTGTAAATGGCCAATTTTAAGACCATCAGAATC GTTTTAGAGCTAGAAATAGC | Cloning |
| 6875 | TGTATTTTCAGGGCGCCATGATGCAATTTAAAGGGGAGAAAA | Cloning |
| 6876 | AGGCCTTTGAATTCCGGATCTTAAGCTTGCGGTTGTGGTT | Cloning |
| 6878 | TGTATTTTCAGGGCGCCATGAACAGTAAATTTCTGTACAATGGG | Cloning |
| 6879 | AGGCCTTTGAATTCCGGATCCTATTTTTTTAATTCTTTATAAGAACTATTTGATAT | Cloning |
| 6886 | CAGGAAGGGCGAATTCTGCATTTTAGCTTTAAAAGCTGTT | Cloning |
| 6887 | GCTTCTTATTTTTATGCTAGAATTTAAATGTAACCTATTACAGG | Cloning |
| 6888 | AGCGTTAACAGATCTGAGCTAAATGAGGTGAAGTTATGGG | Cloning |
| 6889 | CCGGTCAGATCTGCACTGCATTTTTCATCTTAACATTTATACTATC | Cloning |
| 6890 | TAATAGGTTACATTTAAATTCTAGCATAAAAATAAGAAGCCTGC | Cloning |
| 6891 | CCCATAACTTCACCTCATTTAGCTCAGATCTGTTAACGCT | Cloning |
| 6892 | CGATAGTTATGAAGTGAGCTAAGGAGAGTATGGGATGAGTAATGTGAACAAA | Cloning |
| 6893 | TTATTAAAACTTATAGGATCTTAAGCTTGCGGTTGTGGTT | Cloning |
| 6894 | CGATAGTTATGAAGTGAGCTAAGGAGGTGAAGTTATGGGCAGAAGA | Cloning |
| 6895 | TTATTAAAACTTATAGGATCCTATTTTTTTAATTCTTTATAAGAACTATTTG | Cloning |
| 6896 | GAAGATTATGGTGGAGGAGTTGCCCAAGTTTCATCTACG | Cloning |
| 6897 | GATGAAACTTGGGCAACTCCTCCACCATAATCTTCTTCC | Cloning |
| 6898 | CAGCAGGAGGAGGTGTTGCCCAAACATCTTCAACAGTTTA | Cloning |
| 6899 | CTGTTGAAGATGTTTGGGCAACACCTCCTCCTGCTGATTG | Cloning |
| 6906 | AATTAAACTGTAAATGGCCA AGCTATTCAAATATAATCTA GTTTTAGAGCTAGAAATAGC | Cloning |
| 6922 | CGATAGTTATGAAGTGAGCTAAGGAGACCGACTTATGATACAAATTTTGATC | Cloning |
| 6923 | TTATTAAAACTTATAGGATCTTACTGCTCTGCATTTATTTCT | Cloning |
| 6932 | CGATAGTTATGAAGTGAGCTAAGGAGATAATGCTATGAACAGAAAAAGATTGAC | Cloning |

Supplemental Table 4. Oligonucleotides used in this study.

| Oligo | Sequence | Use |
|---|---|---|
| 6933 | TTATTAAAACTTATAGGATCTCATTGGTTCGCCTCCTGAA | Cloning |
| 7013 | CGATAGTTATGAAGTGAGCTCTAAGGAGGGGAATGAAATGCAACAAAATGTAGCAGTA | Cloning |
| 7014 | TTTATTAAAACTTATAGGATCCTTATTTTTTCTTGTACGAGCTTGT | Cloning |
| 7015 | CGATAGTTATGAAGTGAGCTCTAAGGAGGAACGTCATATGGAAGGTAGTCGGGTCCAA | Cloning |
| 7016 | TTATTAAAACTTATAGGATCCTCAGGATGCCTGACTGCTGGC | Cloning |
| 7017 | CGATAGTTATGAAGTGAGCTCTAAGGAGGAGAATAAGATGGCTGCAGGAAGTCAGAGA | Cloning |
| 7018 | TTATTAAAACTTATAGGATCCTTACTGTGCCGGCTGTACCTG | Cloning |
| 7021 | CGATAGTTATGAAGTGAGCTCTAAGGAGGTCAGCTAAATGATTATTGTAATTATTATAAGA | Cloning |
| 7022 | TTATTAAAACTTATAGGATCCTTAAAATACTACATTAGAATCTGATGTTTCTGT | Cloning |
| 7023 | CGATAGTTATGAAGTGAGCTCTAAGGAGGATGATATTTTGCAACTTCTCAAAAAACTC | Cloning |
| 7024 | TTATTAAAACTTATAGGATCCTTACTGTCTTGAATACCAATCGC | Cloning |
| 5465 | AAAATGTTGTAAAAAGACATGAAAAATTTTGAGTTTTTATTGG | Cloning |
| 5464 | ATAAAAACTCAAAATTTTTCATGTCTTTTTACAACATTTTATG | Cloning |
| 6734 | AAATCCTATTCTTGAGAACTCCCATACTTCTCCTTACAAT | Cloning |
| 6735 | ATTGTAAGGAGAAGTATGGGAGTTCTCAAGAATAGGATTT | Cloning |
| 5599 | AAGGGATTTTGAAAGGGGTG | Check *ldt1* deletion |
| 5600 | CTGTCGGTGACTGCCTTCTC | Check *ldt1* deletion |
| 5601 | AGTAGCTGGAGGAGCAGGAT | Check *ldt2* deletion |
| 5602 | CCTGTCAATGTAAATGGGTC | Check *ldt2* deletion |
| 5603 | GCCAAAACTTGGGGAATTGA | Check *ldt3* deletion |
| 5604 | GGTTCTCCACAAGACTGTGG | Check *ldt3* deletion |
| 6808 | GCTGAAGAGGCAAATACTTCTGG | Check *ldt4* deletion |
| 6809 | CCACCATTCTCATAATAAAAAAGGTAATCC | Check *ldt4* deletion |
| 6810 | CGGGGAGCTCTTTGTATAACTATAGATGC | Check *ldt5* deletion |
| 6811 | CCCCATTATTTGCTTTTGTAATTCC | Check *ldt5* deletion |

**References Supplemental Information**

1 Trieu-Cuot, P., Carlier, C., Poyart-Salmeron, C. & Courvalin, P. Shuttle vectors containing a multiple cloning site and a lacZ alpha gene for conjugal transfer of DNA from *Escherichia coli* to gram-positive bacteria. *Gene* **102**, 99-104 (1991). https://doi.org:10.1016/0378-1119(91)90546-n

2 Arends, S. J. & Weiss, D. S. Inhibiting cell division in *Escherichia coli* has little if any effect on gene expression. *Journal of bacteriology* **186**, 880-884 (2004). https://doi.org:10.1128/jb.186.3.880-884.2004

3 Christie, P. J., Korman, R. Z., Zahler, S. A., Adsit, J. C. & Dunny, G. M. Two conjugation systems associated with *Streptococcus faecalis* plasmid pCF10: identification of a conjugative transposon that transfers between *S. faecalis* and *Bacillus subtilis*. *Journal of bacteriology* **169**, 2529-2536 (1987). https://doi.org:10.1128/jb.169.6.2529-2536.1987

4 Youngman, P., Perkins, J. B. & Losick, R. Construction of a cloning site near one end of Tn917 into which foreign DNA may be inserted without affecting transposition in *Bacillus subtilis* or expression of the transposon-borne *erm* gene. *Plasmid* **12**, 1-9 (1984). https://doi.org:10.1016/0147-619x(84)90061-1

5 Stabler, R. A. *et al.* Comparative genome and phenotypic analysis of *Clostridium difficile* 027 strains provides insight into the evolution of a hypervirulent bacterium. *Genome biology* **10**, R102 (2009). https://doi.org:10.1186/gb-2009-10-9-r102

6 Müh, U., Pannullo, A. G., Weiss, D. S. & Ellermeier, C. D. A Xylose-Inducible Expression System and a CRISPR Interference Plasmid for Targeted Knockdown of Gene Expression in *Clostridioides difficile*. *Journal of bacteriology* **201**, e00711-00718 (2019). https://doi.org:10.1128/JB.00711-18

7 Kaus, G. M. *et al.* Lysozyme Resistance in *Clostridioides difficile* Is Dependent on Two Peptidoglycan Deacetylases. *Journal of bacteriology* **202** (2020). https://doi.org:10.1128/JB.00421-20

8 Müh, U., Ellermeier, C. D. & Weiss, D. S. The WalRK Two-Component System Is Essential for Proper Cell Envelope Biogenesis in *Clostridioides difficile*. *Journal of bacteriology*, e0012122 (2022). https://doi.org:10.1128/jb.00121-22

9 Shepherd, T. R. *et al.* The Tiam1 PDZ domain couples to Syndecan1 and promotes cell-matrix adhesion. *Journal of molecular biology* **398**, 730-746 (2010). https://doi.org:10.1016/j.jmb.2010.03.047

10 Fagan, R. P. & Fairweather, N. F. *Clostridium difficile* has two parallel and essential Sec secretion systems. *The Journal of biological chemistry* **286**, 27483-27493 (2011). https://doi.org:10.1074/jbc.M111.263889
